# Spatiotemporal proximity labeling tools to track GlcNAc sugar-modified functional protein hubs during cellular signaling

**DOI:** 10.1101/2022.04.13.488185

**Authors:** Yimin Liu, Zachary M. Nelson, Ali Reda, Charlie Fehl

**Affiliations:** Department of Chemistry, Wayne State University, 5101 Cass Avenue, Detroit, MI, United States

## Abstract

A fundamental mechanism that all eukaryotic cells use to adapt to their environment is dynamic protein modification with monosaccharide sugars. In humans, O-linked N-acetylglucosamine (O-GlcNAc) is rapidly added to and removed from diverse protein sites as a response to fluctuating nutrient levels, stressors, and signaling cues. Two aspects remain challenging for tracking functional O-GlcNAc events with chemical strategies: spatial control over subcellular locations and time control during labeling. The objective of this study was to create intracellular proximity labeling tools to identify functional changes in O-GlcNAc patterns with spatiotemporal control. We developed a labeling strategy based on the TurboID proximity labeling system for rapid protein biotin conjugation that we directed to O-GlcNAc protein modifications inside cells, a set of tools we called “GlycoID.” Localized variants to the nucleus and cytosol, nuc-GlycoID and cyt-GlycoID, labeled O-GlcNAc proteins and their interactomes in subcellular space. Labeling during insulin as well as serum stimulation revealed functional changes in O-GlcNAc proteins as soon as 30 minutes of signaling. We demonstrated using proteomic analysis that the GlycoID strategy captured O-GlcNAcylated “activity hubs” consisting of O-GlcNAc proteins and their associated protein-protein interactions. The ability to follow changes in O-GlcNAc hubs during physiological events like insulin stimulation poises these tools to be used for determining mechanisms of glycobiological cell regulation. Our functional O-GlcNAc datasets in human cells will be a useful resource for O-GlcNAc-driven mechanisms.

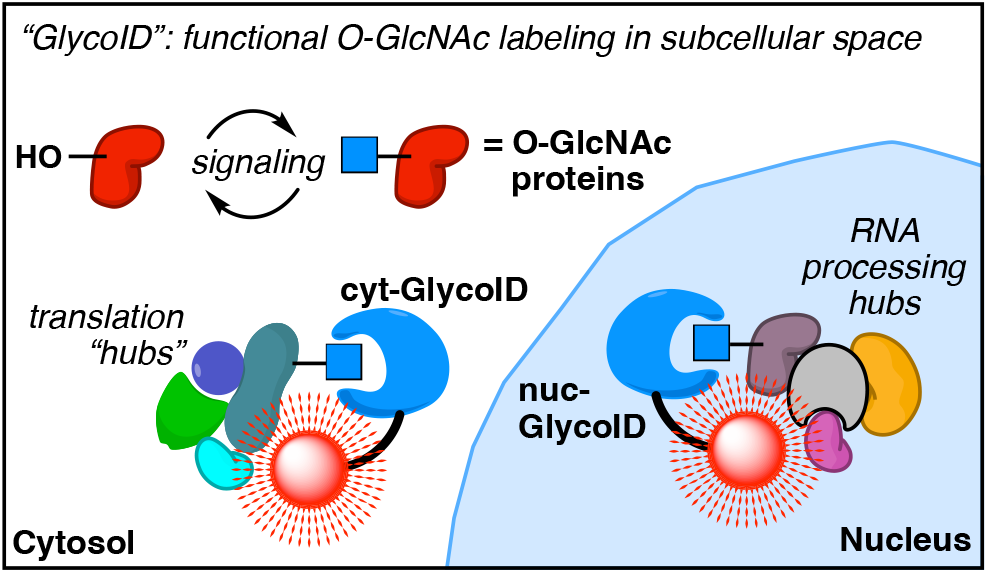

## Introduction

The O-GlcNAc modification on proteins (O-linked N-acetylglucosamine) is a nutrient-and conditions-sensing post-translational modification essential for all mammalian cells to adapt to their microenvironment.^1,2^ Thousands of O-GlcNAc sites^3,4^ regulate cell biology including signaling and transcription in both nutrient-driven^5,6^ and nutrient-independent^7,8^ roles. Protein O-GlcNAcylation is cycled by two proteins, O-GlcNAc transferase (OGT) and O-GlcNAcase (OGA) (**Fig 1A**).^1^ The OGT gene can produce three isoforms, each of which is most active in a distinct cellular location: nucleocytoplasmic ncOGT is primarily found in the nucleus; mitochondrial mOGT is found in mitochondria; and short sOGT, which lacks a nuclear localization signal and therefore is primarily cytosolic.^9,10^ Therefore, a crucial facet of O-GlcNAc regulation depends on the spatial location of target proteins in the cell and which isoform(s) of OGT are produced at a given time (**Fig 1B**).

**Figure 1:**
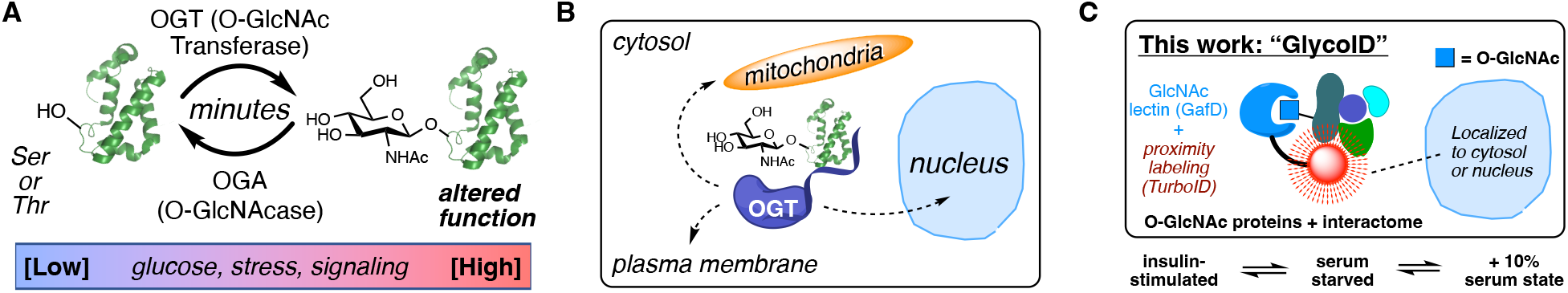
Protein O-GlcNAc post-translational modifications allow cells to rapidly adapt to changes in their environments. **A**) O-GlcNAc modifications are cycled by two enzymes on 1000s of known substrates, occurring at various timescales and driven by fluctuations in nutrients, cell stressors, or signaling cues. **B**) OGT isoforms localize to and move between subcellular compartments, enabling spatiotemporal control of protein functions in cells. **C**) The GlycoID system for live cell O-GlcNAc labeling. The GlcNAc-binding lectin GafD localizes a proximity labeling enzyme TurboID to O-GlcNAcylated proteins. Treatment with the TurboID substrate (biotin) labeled proximal proteins in a ca. 10 nm radius. Subcellular targeting enabled spatial labeling. Changes in cellular O-GlcNAc proteins and proximal interactomes were obtained in live cells under serum or insulin stimulation.

A second mechanism for O-GlcNAc regulation is time-based because O-GlcNAc modifications can be dynamically removed by O-GlcNAcase (OGA). In this vein, mammalian cells regulate the balance of OGT/OGA with respect to overall O-GlcNAc levels, employing a variety of mechanisms including regulatory modifications,^11^ expression,^12,13^ as well as levels of OGT and OGA pre-mRNA transcripts.^14^ In particular, this mRNA regulation via alternative splicing enables cells to respond to O-GlcNAc perturbations within 30 minutes.^14^ During OGT/OGA “rebalancing,” O-GlcNAc events in this 30 minute phase are increasingly recognized as critical for a wide range of cellular functions.^11,15-18^

Recent advances in spatial identification of glycosylated proteins have been developed using proximity labeling tools, including ascorbate peroxidase (APEX)-catalyzed cell-surface galactosamine glycans.^19^ Toward GlcNAc-active enzymes, biotin ligase-catalyzed proximity labeling and identification (BioID) tools have been directed for protein-protein interactomes of OGT^20^ and OGA,^21^ revealing important insights about how each protein is regulated. To-date, however, no proximity labeling tool has been directed toward O-GlcNAc glycan modifications themselves. The lack of an O-GlcNAc intracellular labeling tools makes performing defined spatial and temporal GlcNAc labeling reactions in live cell settings difficult, leading to a recognized gap in strategies to follow O-GlcNAc with spatiotemporal precision.^22,23^

Here, we report a strategy that uses intracellular O-GlcNAc labeling constructs targeted to different locations of the cell. We rely on the recent discovery and optimization of biochemical proximity labeling protein domain, TurboID,^24^ coupled to the O-GlcNAc binding domain GafD^25,26^ to enable intracellular labeling of O-GlcNAcylated proteins and any associated protein complexes. These tools, which we call “GlycoID,” revealed functional O-GlcNAc “hubs” that responded to conditions in live cells including insulin stimulation and serum feeding (**Fig 1C**). Targeted GlycoID constructs to the nucleus or cytoplasm revealed location-specific O-GlcNAc hubs. Additionally, rapid GlycoID labeling conditions within 30 minutes of signal induction confirmed the possibility for tracking O-GlcNAc modifications in real time. Here, we used these tools to expand our view of O-GlcNAc-associated functions during signaling and nutrient sensing.

### Results and Discussion – Design of GlycoID tools for intracellular O-GlcNAc labeling

We designed proximity labeling systems for intracellular, O-GlcNAc-driven protein tagging by fusing the proximity labeling domain miniTurboID (mTurbo) to the GlcNAc-binding lectin GafD^27^ (**Fig 2A**). We chose mTurbo because it uses the non-toxic substrate biotin to attach nonhydrolyzable biotin tags to proteins a tight < 10 nm radius from a bound target protein. The small 28 kDa size of mTurbo enables efficient expression in human cell lines as a fusion to protein targeting domains. Furthermore, since the parent biotin ligase BirA has relatively high substrate concentration requirements (*K*M ≈ 5 µM),^28^ we can cleanly conduct cellular labeling media that lacks biotin, including the popular Dulbecco’s Modified Eagle’s Media. GafD was chosen as the GlcNAc-binding partner of interest because of its selectivity for GlcNAc-linked molecules over other sugars, including >10 fold binding selectivity over glucose-linked molecules.^29^ Additionally, GafD-based fluorescent reporters for OGT activity have been successfully expressed and used in cell studies.^26,30^

**Figure 2:**
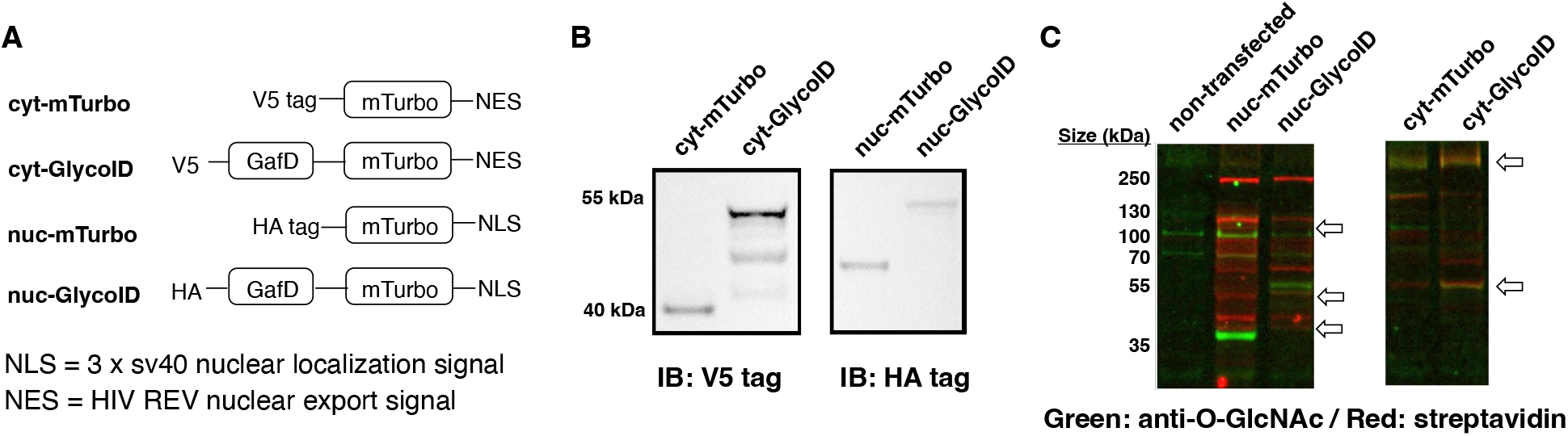
Confirmation of GlycoID labeling. A) Construct design for variant proximity labeling systems. mTurbo = mini-TurboID system only (undirected control). GlycoID = GlcNAc-directed system. cyt = cytosol, nuc = nucleus targeting. B) Expression of the indicated construct in HeLa cells. C) Biotin induction at 100 mM, 6 h reaction followed by cell lysis and total cell blotting. Dual-fluorescence blot was probed with a pan-O-GlcNAc antibody (O-GlcNAc MultiMAb) and visualized with an AlexaFluor-555 secondary (green) and streptavidin-Cyanine5 (red) fluorescent conjugates. Strongly overlapping (gold color) bands are indicated by arrows.

We used the reported structures of GafD (1oio)^31^ and BirA (3rux)^32^ to design our constructs bearing an N-terminal GafD, a short linker containing an HA or V5 tag for immunoblotting, and the C-terminal mTurbo domain containing a localization sequence (full sequences found in **Supplementary Table 3**). We hypothesized that mTurbo-GafD constructs, hereafter referred to as “GlycoID,” would bind GlcNAc residues on intracellular proteins for covalent biotin labeling. Because TurboID constructs label proteins in a 10 nm radius around their binding partners, we additionally expected that GlycoID could label the interactomes of O-GlcNAcylated proteins *in situ*. Transient binding/release of the constructs from targets could thereby label a diverse array of O-GlcNAcylated protein complexes during intracellular functions, including signaling events such as insulin stimulation.

O-GlcNAcylation primarily occurs in the nucleus and cytoplasm of cells.^33^ We generated **cyt-GlycoID** and **nuc-GlycoID** using mTurbo plasmids contributed by the Ting group (**Fig 2A**).^24^ As control constructs that were not directed to GlcNAc-modified proteins, we expressed cyt-mTurbo and nuc-mTurbo, constructs that lacked the GafD domain. Transient expression of all four constructs proceeds smoothly in HeLa (**Fig 2B**) and HEK293T cells (**Supplementary Figure 1**). The activity of these constructs to biotinylate intracellular proteins was shown by incubating GlycoID-expressing cells with 100 mM biotin for 6 h, followed by cell lysis and immunoblotting for O-GlcNAcylation and biotinylation (**Fig 2C**). Confirmation of O-GlcNAc binding and labeling activity was shown by dual imaging with anti-O-GlcNAc/AF555 (**Fig 2C, green signals**) overlaid with streptavidin/Cy5 (**Fig 2C, red signals**). Gold overlapping bands suggested on-target GlycoID labeling of O-GlcNAcylated proteins in HeLa cells.

### Optimization and validation of intracellular O-GlcNAc labeling

We characterized the labeling activity of GlycoID constructs following transient expression. To initiate proximity labeling, we expressed GlycoID constructs in HeLa cells for 48 h. Treatment of biotin in a range from 25-500 mM concentrations was applied from an aqueous stock, and labeling reactions were allowed to proceed from 10 min to overnight (**Fig 3A**). We observed significant, dose and time-dependent labeling of soluble nucleocytoplasmic proteins in HeLa cells. Balancing concentration and time identified detectable labeling as soon as 10 min at 500 mM [biotin], compared to maximal labeling which peaked under all concentrations at 6 h. Overnight labeling did not lead to stronger labeling. Full activity blots are shown in **Supplementary Figure 2**.

**Figure 3:**
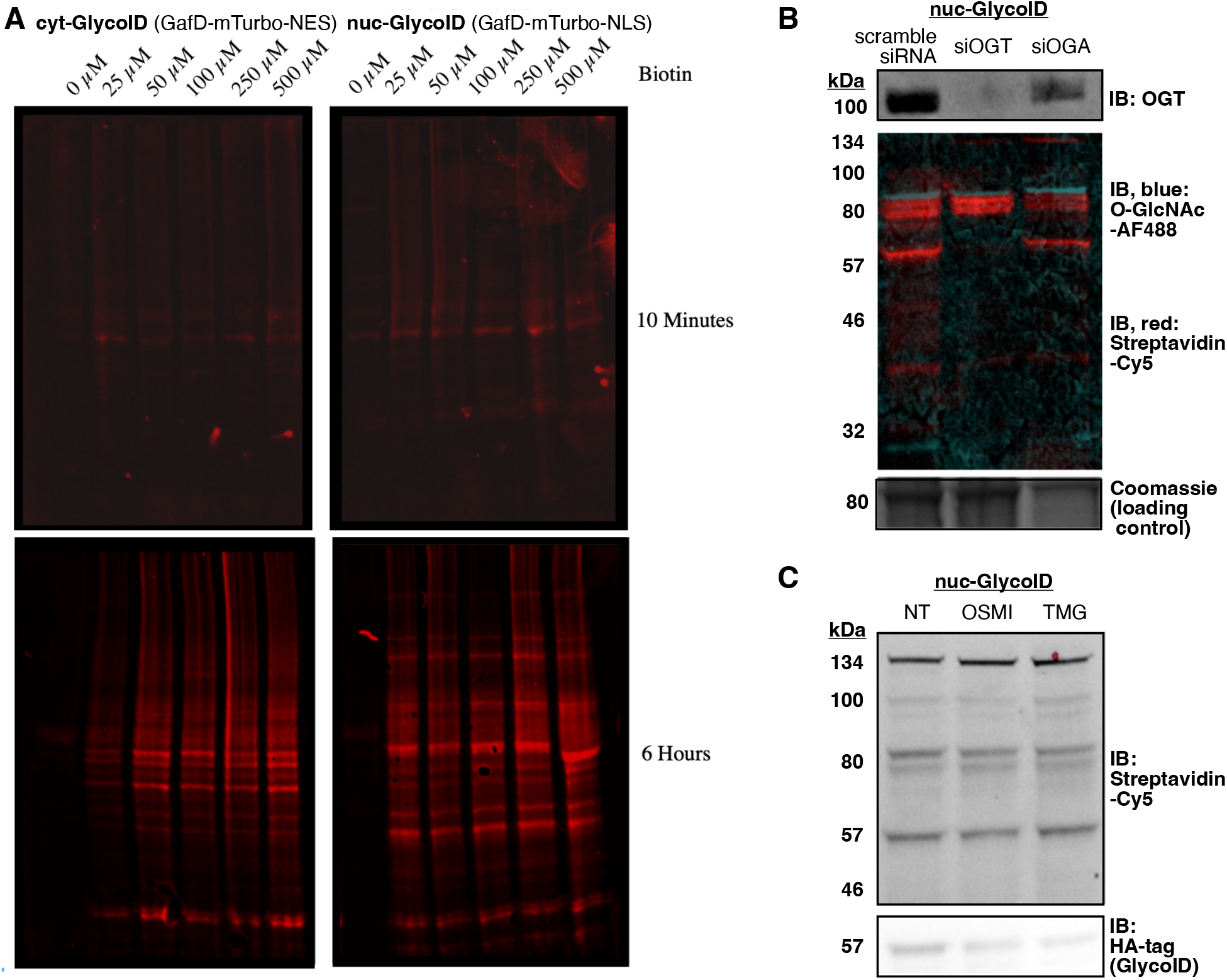
Activity optimization for GlycoID labeling in HeLa cells. **A**) Concentration and time course for the indicated construct. Full blots shown in **Supplemental Figure 2**. Biotinylated protiens were visualized using a streptavidin-Cy5 fluorescent conjugate. **B**) O-GlcNAc engineering with RNA silencing of OGT or OGA (siOGT or siOGA, respectively) revealed altered GlycoID labeling. Knockdowns were initiated 24 h prior to transfection with GlycoID constructs and maintained during labeling. **C**) Inhibition studies using OSMI-4 (OGT inhibitor, 40 µM) and thiamet-G (TMG, OGA inhibitor, 10 µM) and subsequent GlycoID labeling.

We engineered overall GlcNAc levels in cells to confirm specificity for GlcNAc-driven protein labeling. Validation of O-GlcNAc binding by our constructs came from control experiments where either OGT or OGA, the two enzymes that add and remove O-GlcNAc, were suppressed before nuc-GlycoID labeling was induced with biotin treatment. We found that knockdown of OGT led to a suppression of OGT protein levels, total O-GlcNAc levels, and the number of biotin-labeled protein bands after 24 h of OGT RNA silencing (siRNA) treatment (**Fig 3B**). On the other hand, knockdown of OGA was expected to elevate both O-GlcNAc levels and labeling. We observed similar O-GlcNAcylation levels and number of biotin bands vs. scrambled siRNA control (**Fig 3B, left vs. middle**), but noted that OGT levels were also slightly diminished upon OGA knockdown. Tight regulation of OGT and OGA levels have been reported,^13,14^ so we propose that our OGA knockdown studies led to concomitant, feedback-based reduction of OGT to compensate elevated O-GlcNAc levels via OGA loss. This paradoxical response of OGT upon OGA knockdown may explain the lack of appreciable difference with the scramble siRNA control.

Inhibition of OGT with the small molecule OSMI-4 prior to biotin induction led to roughly similar levels of nuc-GlycoID labeling (**Fig 3C, middle**). However, pre-treatment of cells with the OGA inhibitor thiamet-G (TMG) led to slightly increased labeling and number of bands relative to total protein loading (**Fig 3C, right**). Full blots for the GlcNAc engineering experiments are shown in **Supplementary Figure 3**. Overall, the O-GlcNAc suppression and elevation results had a subtle effect on GlycoID-driven labeling patterns but followed the expected trends. Mammalian cells are well established to rapidly attenuate OGT and OGA protein levels upon inhibition or knockdown of either species in order to maintain O-GlcNAc homeostasis, so we hypothesize that OGT/OGA regulation over the 24 h periods used for O-GlcNAc elevation or suppression led to reduced impact on overall GlycoID labeling patterns.^14^

### GlycoID identifies physical hubs of O-GlcNAcylated protein clusters in subcellular space

The targeted constructs cyt-GlycoID vs. nuc-GlycoID gave a striking difference in band patterns (**Fig 3A**), as we expected from labeling between different subcellular locations. To compare specific proteins between the two cellular compartments, we carried out quantitative tandem mass spectrometry-based proteomics (LC-MS/MS) to identify which proteins are labeled by our targeted GlycoID constructs. HeLa cells that expressed nuc-GlycoID were compared with cells that expressed nuc-mTurbo as a non-sugar directed control in replicates of 4 per condition. To prevent premature labeling each construct was expressed in DMEM media, which completely lacks biotin (see **Fig 3A**, 0 µM biotin). Biotin labeling was induced at 48 h post-expression by adding 100 µM biotin and incubating for 6 h at 37 °C to obtain maximum intracellular labeling. High-confidence proteomic hits were chosen based on significance (p < 0.05) and fold-enrichment (log2 > 0.5) cutoffs using Perseus version 1.6.2.1. Hits also had to satisfy the condition of being detected in at least 3 of the 4 replicates and to have at least 3 unique peptide matches during MaxQuant processing. Blots that confirmed similar labeling efficiency between all replicates are shown in **Supplementary Figure 4**.

In the nuclear-targeted experiment, we identified 98 high-confidence proteins exclusive to nuc-GlycoID (vs. nuc-mTurbo) (**Fig 4A**). Volcano plot analysis showed an additional 4 proteins enriched by nuc-GlycoID over nuc-mTurbo (**Fig 4B**). Comparison with the O-GlcNAcome^33^ databank revealed that 49% of these 102 enriched/exclusive proteins were known O-GlcNAc proteins. A unique strength of proximity labeling is revealing proteins that physically associate with the target proteins in subcellular space.^34^ To identify whether the remaining 51% of these proteins correspond to O-GlcNAc protein binding partners, we used the STRING-db^35^ to analyze reported protein-protein interactions (PPIs) between the nuc-GlycoID dataset. We set STRING to only utilize experimentally verified PPIs. Interestingly, the majority of the non-O-GlcNAc protein hits (31/52) are known to physically associate with O-GlcNAc proteins labeled by nuc-GlycoID. Based on the labeling radius of TurboID,^24^ these proteins were expected to be within 10 nm of the target bound GlcNAc glycoproteins. Gene ontology analysis using k-means clustering revealed that these associated proteins comprised five “functional O-GlcNAc hubs”: mRNA binding; transcription factors; nucleotide binding; gene expression; and splicing (**Fig 4C**). The PPI enrichment p-value, which compares the observed connections (edges between nodes) in the interaction network against what would be expected with no enrichment, averaged between p = 1.19e-05 to <1.0e-16. For example, the highly-connected splicing cluster had 60 observed PPI edges in our STRING analysis vs. the expected number of 5 “random” PPIs based on the sizes of these proteins alone. This extremely high enrichment of functional protein “hubs” reveals that nuc-GlycoID labeled known interaction partners of O-GlcNAcylated proteins with very high confidence (significance). A summary of the clustering analysis with key O-GlcNAc proteins identified is shown in **Table 1**.

**Table 1:** GlycoID labels O-GlcNAc clusters in defined cell locations. Full activity conditions: 6 h with 100 µM biotin.

|  | Cluster | # of nodes (proteins) | # of edges (interactions) | PPI enrichment p-value | Sample proteins |
| --- | --- | --- | --- | --- | --- |
| <u>nuc-GlycoID</u> | mRNA binding | 28 | 46 | <1.0e-16 | DHX9, RMB14, ZFR, WBP11 |
|  | splicing | 19 | 60 | <1.0e-16 | SF1, SF3A1, SF3B2, RBM25 |
|  | transcription factors | 19 | 5 | 4.59e-05 | HCFC1, JunB, CCAR2, FUBP1 |
|  | gene expression | 19 | 14 | 1.19e-05 | SYMPK, MBNL1, NONO, COIL, AGFG1 |
|  | Metabolism and signaling | 14 | 7 | 5.41e-05 | NOLC1, DHDH4, DIDO1, NKRF |
| Nuc-mTurbo | None | 14 | 1 | 1.0 (no significance) | HSPA8, NUMA, HIST1H1D |
| <u>cyt-GlycoID</u> | translation | 26 | 31 | 2.39e-10 | EF1A1, RPL18, LYAR, LLPH |
|  | RNA binding | 24 | 10 | 3.34e-06 | GNL3, RRP1B, THOC2 |
|  | cytoskeleton | 17 | 16 | 6.66e-16 | ACTB, ACTA, RRBP1 |
| <u>cyt-mTurbo</u> | translation | 22 | 48 | 4.44e-16 | RPL3, EIF5B, RPS26 |
|  | glycolysis | 7 | 6 | 2.217e-11 | GAPDH, ALDOA, ENO1 |
|  | ubiquitinylation | 9 | 2 | 0.00226 | UBAP2, TRIP12 |
|  | spliceosome | 12 | 2 | 0.0337 | SRSF11, SREK1, U2SURP |

**Figure 4:**
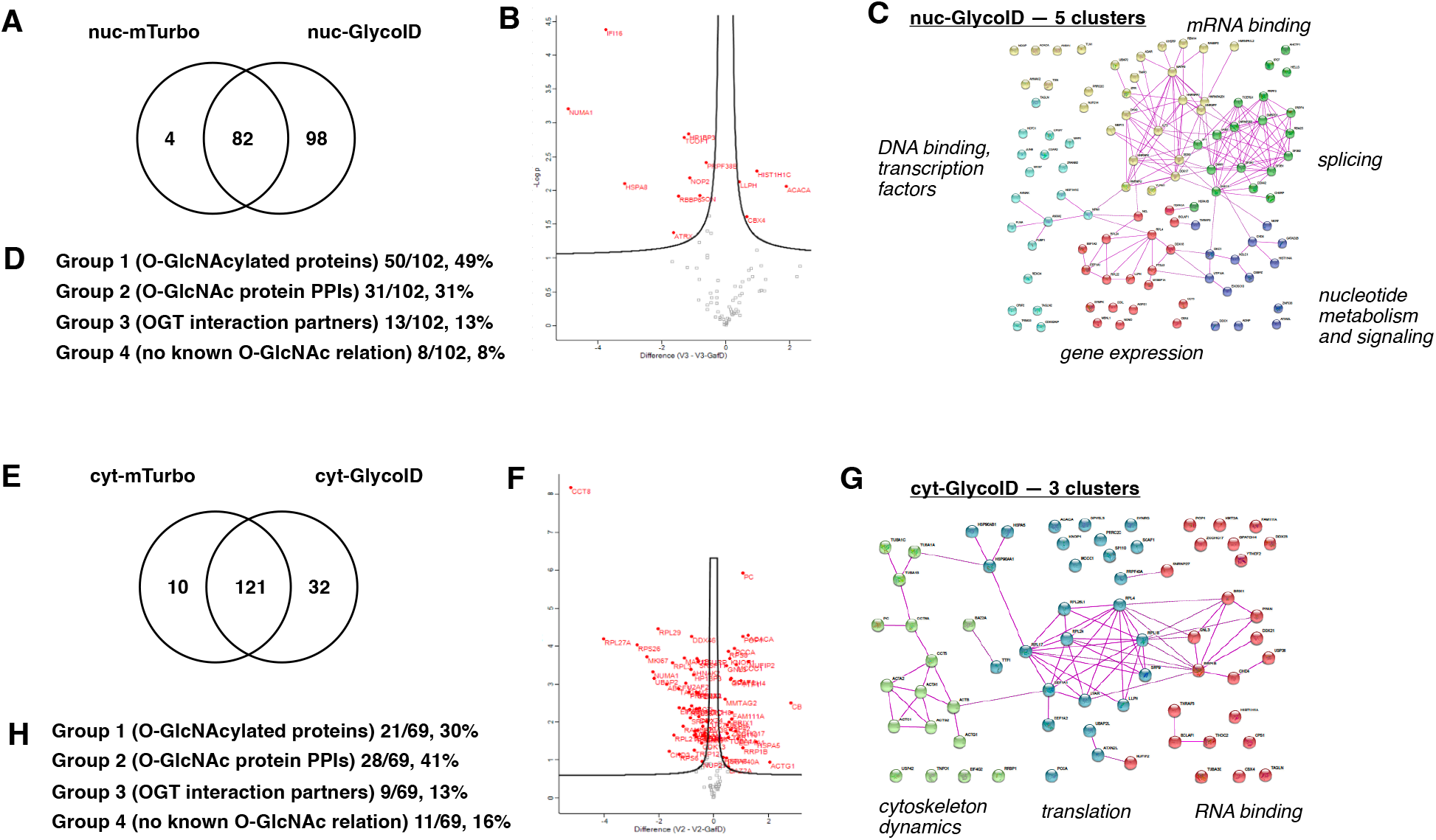
Quantitative proteomics with GlycoID constructs. **A**) Exclusive hits between GlcNAc-binding nuc-GlycoID and control nuc-mTurbo. **B**) Enrichment analysis between nuc-mTurbo and nuc-GlycoID, statistically significant hits are shown above the volcano plot. **C**) Physical interactions between nuc-GlycoID hits reveal functional clusters with key O-GlcNAc linkages. **D**) The protein groups labeled by nuc-GlycoID, as defined in text. **E**)-**H**) Analysis for cyt-GlycoID vs. cyt-mTurbo. Full-sized STRING plots with labeled O-GlcNAc hits are found in the **Supporting Figures 5** and **7-9**.

Further analysis of possible sources of O-GlcNAc-driven labeling on the remaining proteins without known O-GlcNAc sites or PPIs with O-GlcNAcylated proteins was performed using OGT-PIN, the O-GlcNAc transferase protein-interaction network.^36^ OGT is known to form functional complexes with various activity hubs in cells,^37^ including histone chaperone complexes^38^ and tet protein DNA demethylation complexes,^39^ and there is a strong likelihood of coincidental proximity-based labeling of proteins within 10 nm radius of an OGT hub. We used the OGT-PIN to reveal that a third group of proteins labeled by GlycoID constructs overlap with OGT-PIN data. Together, we divided the GlycoID analysis into four groups: Group 1 proteins with known O-GlcNAc sites; Group 2 proteins that interact with O-GlcNAcylated proteins (via STRING-db); Group 3 proteins that complex with OGT; and Group 4 proteins with no O-GlcNAc connection, likely experimental noise from high abundance proteins like thioredoxin. The nuc-GlycoID results from HeLa cells are summarized in **Fig 4D**. The full lists of gene names, statistics, and fold-enrichment values for all identified proteins are found in the **Appendix Table 1**. The full STRING plot, with labeled groups, is found in **Supplementary Figure 5**.

We also performed STRING analysis on the non-targeted nuc-mTurbo-only constructs (**Supplementary Figure 6**). Only 14 proteins in total were unique or enriched by the nuc-mTurbo construct. These proteins did not cluster into any discrete functions (**Table 1**) and were mostly high abundance proteins like heat shock proteins (HSPA8), histones (HIST1H1D), and microtubule binding proteins (NUMA1), which indicated that nuclear-mTurboID labeling was dictated more by protein abundance. The full STRING plot, with labeled groups, is found in **Supplementary Figure 7**.

Next, we conducted the cytosolic GlycoID experiment by comparing cyt-GlycoID labeling with cyt-mTurbo-expressing HeLa cells. After a 6 h induction with 100 µM biotin, 32 high-confidence proteins were exclusive to the cyt-GlycoID (**Fig 4e**). Volcano plot analysis revealed an additional 37 proteins significantly enriched between cyt-GlycoID and cyt-mTurbo (**Fig 4f**). STRING-db analysis of PPIs reveal that cyt-GlycoID is also able to identify functional O-GlcNAc hubs in the cytosol of cells (**Fig 4g**). For cyt-GlycoID, 30% of the dataset (21 hits) were known O-GlcNAc proteins, and the majority of the non-O-GlcNAc labeled hits (28 of the remaining 49) are known to be physically associated with these O-GlcNAc proteins (**Fig 4h**). The cyt-GlycoID results in HeLa cells gave three major clusters: RNA binding; cytoskeleton dynamics; and translation. The PPI enrichment p-values ranged from p = 3.34e-6 to 6.66e-16, revealing that cyt-GlycoID labels known protein clusters with high statistical significance. A summary of cyt-GlycoID clusters and key O-GlcNAc proteins is found in **Table 1**. The full STRING plot, with labeled groups, is found in **Supplementary Figure 8**.

Conversely, we observed four significant functional clusters observed with cyt-mTurbo (**Supplementary Figure 6**). The cyt-mTurbo functional clusters had diverging roles of ubiquitinylation, glycolysis, and spliceosome (not observed in cyt-GlycoID) and one overlapping role, translation (**Table 1**). These different labeled functions indicates that cyt-mTurbo and cyt-GlycoID were directed to different labeling complexes over the 6 h labeling period. The full STRING plot for cyt-mTurbo with labeled groups is found in **Supplementary Figure 9**.

Statistical analysis was performed on these four datasets to determine whether these GlycoID tools label known O-GlcNAcylated proteins more frequently than random proteins. We used a Fisher’s exact test analysis of the targeted (GafD-mTurbo, GlycoID) and non-targeted (mTurbo-only) constructs to test the null hypothesis that GlycoID labels O-GlcNAc proteins with equal frequency than random proteins (see **Supplemental Discussion**). The probability for higher GlcNAc labeling was above the threshold to reject the null hypothesis (**Supplementary Table 7**). Hoever, our data revealed that a significant number of proteins were physically associated with known O-GlcNAcylated proteins. When we incorporated both Group 1 (known O-GlcNAcylated proteins) + Group 2 (known interactome of O-GlcNAcylated proteins), our Fisher’s exact tests achieved significance for both cyt-GlycoID (p = 0.001) and nuc-GlycoID (p = 0.034) (**Supplementary Table 8**). This analysis indicated that targeted GlycoID tools enriched O-GlcNAcylated proteins with their known interactomes at high confidence levels.

Among the directly O-GlcNAcylated proteins we observed with nuc-GlycoID, HCFC1, JunB, SF1, and ZFR stood out because they are among the top 10% of the O-GlcNAcome, based on the “O-GlcNAc score” from 0-100 that ranks the strength of the evidence for an O-GlcNAc site on a given protein.^33^ These nuclear proteins are most involved in transcriptional regulation and in production and splicing of mRNA, dynamic nuclear functions that O-GlcNAc is known to regulate.^14,40^ Among the cyt-GlycoID hits, EF1A1, ACTB, and RRBP1 have high O-GlcNAc scores and are involved translation and cytoskeletal movements, two key cytosolic functions regulated by O-GlcNAcylation.^41,42^

One of the distinctive features of using an O-GlcNAc-targeted proximity labeling system is the ability to observe O-GlcNAcylated “functional hubs” made up of protein-protein interactions. The extremely high number of PPIs (up to 60) and the strong p-values between 10^−5^ and < 10^−16^ suggested that the GlycoID strategy not only identified O-GlcNAcylated proteins but also their key interaction partners. These O-GlcNAc interactomes may also be proximally involved in O-GlcNAc regulated functions. These major clusters focus on transcription and mRNA splicing in the nucleus and translation in the cytosol, which is consistent with known O-GlcNAc transferase roles in mammalian cell proliferation^37^ and nutrient sensing.^43-45^

### Functional O-GlcNAc glycoproteomics of nutrient sensing and insulin signaling

The intracellular nature of GlycoID allows monitoring of O-GlcNAc events in real-time and in localized subcellular space. We used our proteomic analysis workflow to analyze O-GlcNAc-related functions in cells to compare the effects of insulin stimulation following overnight serum starvation. Insulin is known to trigger changes in OGT and O-GlcNAcylation levels.^11,46,47^ Furthermore, engagement of the insulin receptor causes a rapid change in OGT localization from the nucleus to the plasma membrane and cytosol between 5-30 min.^11^ After 60 min, OGT leaves the plasma membrane and returns to the nucleus.^11^ Therefore, the spatiotemporal features of the GlycoID strategy were poised to track functional effects of O-GlcNAc during insulin signaling.

We hypothesized that nuc-GlycoID and cyt-GlycoID could detect key changes in O-GlcNAc-driven functional “hubs” following starvation vs. stimulation. For these experiments, we reduced the labeling time to 30 minutes to fall within the known time that insulin is known to trigger changes in OGT activity.^11^ In our initial characterization, we observed reliable labeling at short time points at higher biotin concentration, so we raised the biotin in these experiments to 500 µM for these 30 minute labeling reactions. We performed four replicates of each condition, and hits chosen were observed in at least 3 of 4 for the analysis for nuc-GlycoID starved, + serum, or + insulin and cyt-GlycoID starved, +serum, or + insulin. For the starved cyt-GlycoID version, 2/4 proteomic runs failed to give quality datasets. For this condition alone (cyt-GlycoID, starved), we chose hits that were observed in both successful proteomic replicates. Statistical validation was performed in Perseus, cutoffs of p = 0.05, fold-change > +/- 0.5, and at least 3 unique peptide matches for a protein to be assigned as a high-confidence hit for further analysis. Full protein blots that confirmed labeling efficiency between all replicates are show in **Supplementary Figure 10**. The full lists of gene names, statistics, and fold-enrichment values for all identified proteins in the starved vs. stimulated cell conditions are found in the **Appendix Table 2**.

We used nuc-GlycoID to compare O-GlcNAc proteins between serum starved and insulin-stimulated O-GlcNAc-related proteins in the nucleus. We observed changes between the nucleus, where 19 proteins were enriched/exclusive between the “starved” cells and 22 were enriched between the “insulin-stimulated” cells (**Fig 5A/B**). Of the enriched/exclusive proteins in the nuc-GlycoID insulin dataset, a relatively low 32% were O-GlcNAc modified. We used STRING-analysis to identify reported protein interactions between the hits (**Fig 5C**). We also used the OGT-PIN set to analyze the OGT interactors and split the analysis into the O-GlcNAc-related Groups 1-4, as detailed above (**Fig 5D**). Our k-means clustering analysis in STRING revealed only one significant functional group, involved in pre-ribosomal assembly. The remaining proteins did not fall into a significant grouping, but included O-GlcNAcylated regulatory proteins like ABCF1 (translation initiation) and CDK12, a kinase involved in regulating the cell cycle.^48^ The k-means clustering results are summarized in **Table 2**. The full STRING plot, with labeled groups, is found in **Supporting Figure 11**.

**Table 2:**
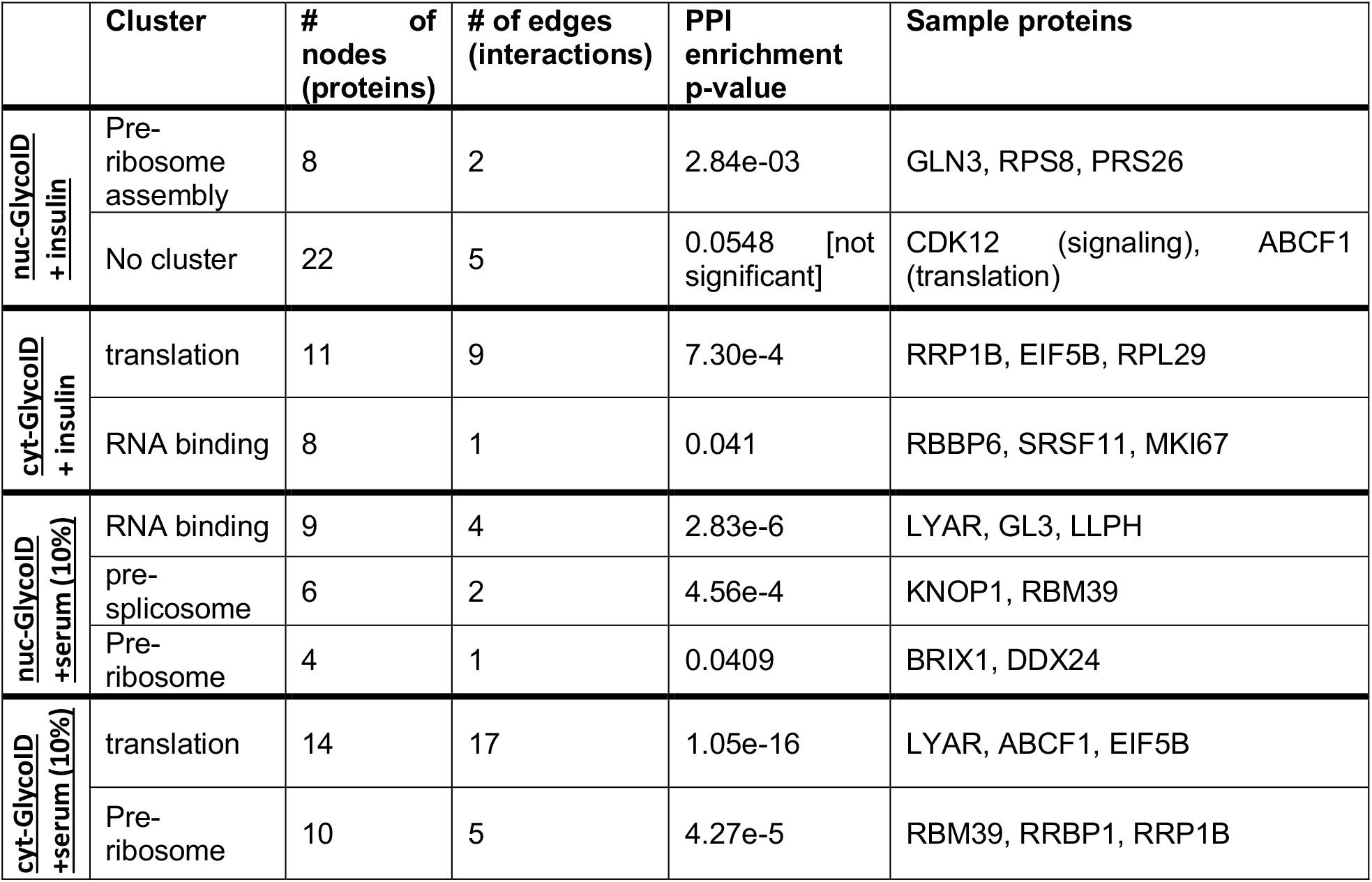
Functional O-GlcNAc labeling in stimulated vs. serum starved cells. Labeling: 30 min with 500 µM biotin.

**Figure 5:**
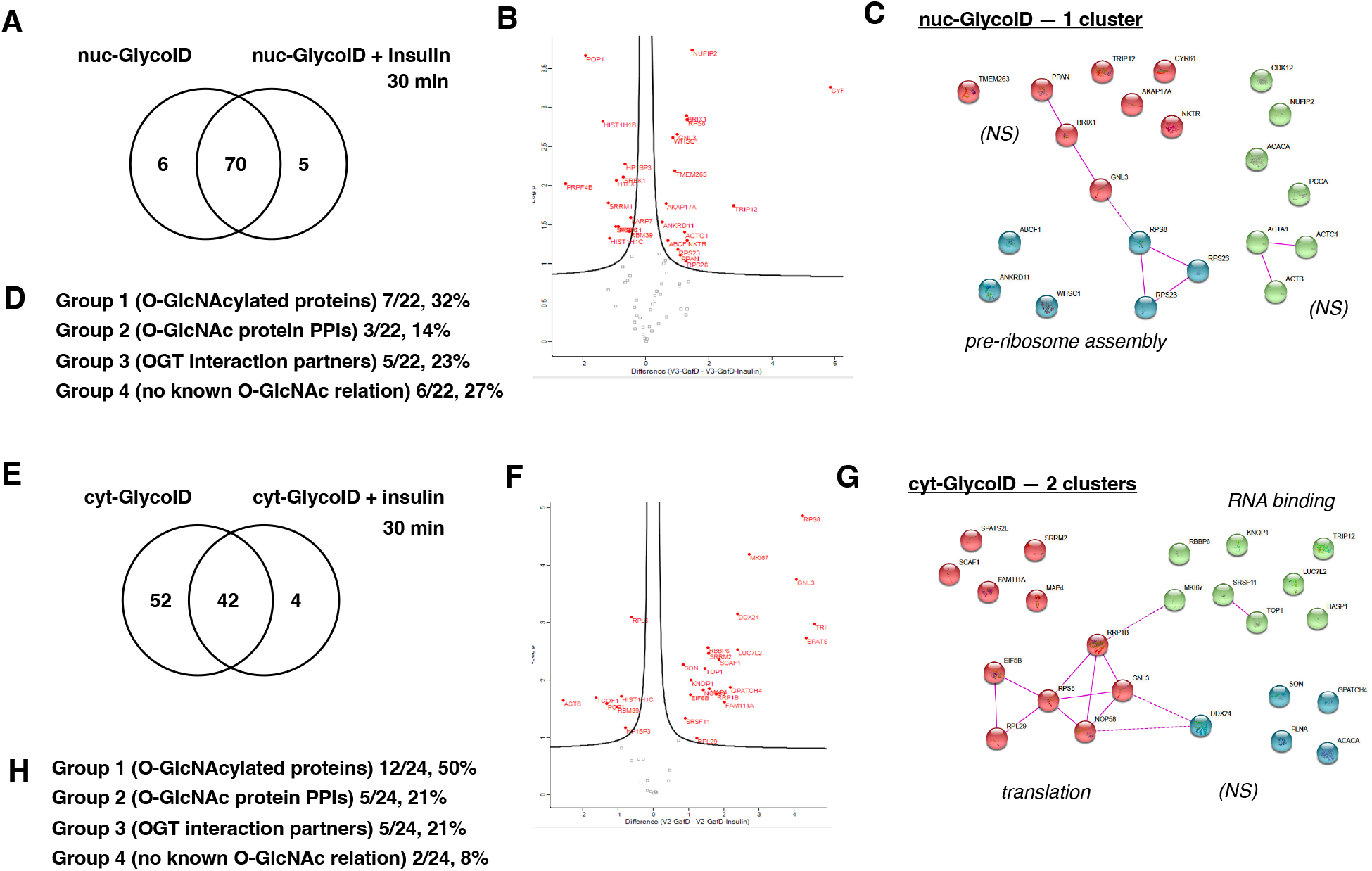
Functional O-GlcNAc proteomics with GlycoID during insulin signaling. a) Exclusive hits between serum-starved nuc-GlycoID and insulin-stimulated nuc-GlycoID, 30 min at 500 µM biotin labeling.**B**) Enrichment analysis between starved and insulin conditions, statistically significant hits are shown above the volcano plot. **C**) Physical interactions between nuc-GlycoID/insulin hits revealed one functional cluster with key O-GlcNAc linkages. **D**) The protein groups labeled by nuc-GlycoID/insulin, as defined in text. **E**)-**H**) Similar analysis for cyt-GlycoID-starved vs. cyt-GlycoID + insulin. *Fully sized STRING plots, with labeled O-GlcNAc hits, are found in* ***Supporting Figures 11-12***.

The cytosolic analysis revealed 65 starved hits vs. 24 insulin-driven proteins (**Fig 5E/F**). STRING analysis and k-means clustering identified two significant clusters, primarily involved in translation and RNA binding (**Fig 5G**). There were also several non-clustered proteins with involvement in actin dynamics (FNLA) and fatty acid biogenesis (ACACA), two features that could respond to insulin. The Group 1-4 analysis revealed that, for known O-GlcNAc proteins, cyt-GlycoID (50% Group 1 labeling) worked significantly better than nuc-GlycoID (32% Group 1 labeling) under insulin, 30 minute stimulation (**Fig 5H**). The functional k-means clustering results are summarized in **Table 2**. The full STRING plot, with labeled groups, is found in **Supporting Figure 12**.

We also used the serum-starved cells to compare with serum-fed conditions, again at the 30-minute time point with 500 µM biotin. Because serum contains both nutrients and growth factors, we hypothesized that GlycoID labeling patterns from cells stimulated by serum would also display different functional hubs. We conducted proteomic analysis as for the insulin labeling (**Supplementary Figure 13**). We observed 3 significant clusters in the nuc-GlycoID + serum dataset, RNA binding, pre-spliceosome assembly, and pre-ribosome assembly (full STRING plot in **Supplementary Figure 14**). In the cyt-GlycoID + serum dataset, we observed two significant clusters: translation and pre-ribosome assembly. The full STRING plots, including groups, are found in **Supplemental Figures 14**-**15**.

Overall, we observed fewer total exclusive and enriched proteins in both functional experiments compared to the first analysis of mTurbo vs. GlycoID constructs. We note that in these functional experiments we compared GlycoID with or without insulin and serum, and therefore we lost many overlapping O-GlcNAcylated proteins that do not change O-GlcNAc status during stimulation that were labeled by the active GlycoID construct in both conditions. This is expected due to the overall widespread distribution of O-GlcNAc on proteins and suggested that only a subset of O-GlcNAcylated proteins actively responded to insulin or serum stimulation at the 30-min time point. Longer induction times might reveal more widespread changes in O-GlcNAc patterns and interactomes. In this insulin labeling experiment, we observed diminished nuc-GlycoID labeling and enhanced cyt-GlycoID labeling, which is approximately the inverse of what we observed under steady-state cell conditions, where OGT is more active in the nucleus (compare **Fig 4D-H** vs. **Fig 5D-H**). This spatiotemporal effect might reflect some movement of OGT toward the plasma membrane that is reported to peak at 30 minutes.^11^ Though we have far from completed a full time course of insulin signaling and O-GlcNAc events, which is known to last until at least 4 h post-stimulation, our GlycoID tools are poised to track O-GlcNAc dynamics in live cells over a variety of homeostasis, signaling, and pathological conditions.

## Conclusions

Intracellular O-GlcNAc patterns change in subcellular space and in real-time during signaling processes, but tools to capture these functional changes in live cells remain an emerging area. We demonstrate that proximity labeling protein constructs targeted to GlcNAc modifications— GlycoID—is an intracellular strategy label O-GlcNAcylated proteins and their interactomes. Spatial targeting using localization signals revealed different labeling patterns between O-GlcNAc interactomes in the nucleus vs. cytosol. Furthermore, functional O-GlcNAc labeling experiments conducted for short, 30-min periods during insulin-or serum-stimulation demonstrated the ability to track O-GlcNAcylation patterns and interactome changes in real-time. This functional O-GlcNAc interactome data adds evidence to a growing area of OGT-regulated “hubs” of activity,^36^ including splicing,^14^ metabolism,^37^and signaling.^48^ Despite these advances, we note limitations to our current study that we will be addressing. Here, all experiments and data reported in this paper were performed with transient expression in human cell lines. We felt that this would be the most useful sharable tool, and our constructs will be available on the Addgene databank. Further applications are possible with cell lines stably-expression GlycoID constructs. A more detailed time courses of insulin signaling using stable-GlycoID constructs is underway in our laboratory. Improved control over labeling can also be envisioned with peroxidase-based reagents,^19^ though we initially chose to avoid APEX strategies due to the requirement for hydrogen peroxide treatment, a factor reported to modulate O-GlcNAc patterns.^21^ Future efforts will also be directed toward disease models, where genetic engineering can be used to further validate GlycoID labeling outcomes. At present, our tools and functional datasets can serve as useful resources for determining functional effects of dynamic O-GlcNAc modifications in live cell settings.

## Methods

### In vitro proximity labeling of live cells

HeLa or HEK293T cells were seeded on sterile plates and transfected with cyt-mTurbo, nuc-mTurbo, cyt-GlycoID, or nuc-GlycoID with at least 48 hours of incubation in DMEM (Dulbecco’s Modified Eagle Medium) at 37 °C. Media was then replaced with media supplemented with 100 µM biotin and allowed to incubate for 6 hours. Then cells were rinsed with phosphate-buffered saline (PBS) twice before freezing. Cells were harvested via scraping cells off the plates with RIPA buffer, lysing the cells with sonication. Inhibition studies were performed with Thiamet-G (10 µM), OSMI-4 (40 µM), and with siRNA knockdowns of OGA and OGT. For chemical inhibition, post-transfected cells are incubated with the appropriate compound at least 24 hours before biotin labeling. For siRNA KD, cells were first transfected with Dharmacon™ ON-TARGET plus SMART pool human OGT siRNA, SMART pool human OGA siRNA, or ON-TARGET control pool non-targeting pool siRNA 24 hours before transfection with GlycoID tools, followed by 48-72 hours of incubation before biotin addition. Plasmids generated in this research are available via the Addgene repository with ID# 184640 (cyt-GafD-mTurboID-V5) and ID# 184641 (nuc-GafD-mTurboID-HA).

### Preparation of Samples for Proteomics

Following biotin labeling, cells were harvested with 100 µL RIPA buffer (150 mM NaCl, 0.5 mM tris, 1% NP40, 0.1% SDS) and were lysed via passage through needle (at least 10 passes). To enrich biotinylated proteins, samples containing 400 µg total protein were incubated with streptavidin-coated magnetic beads overnight at 4 ^0^C. The beads were then washed with RIPA buffer, wash buffer (50 mM Tris, pH 7.4, 2% SDS), and twice more with RIPA buffer. The beads were then resuspended in DTT in PBS and were then treated with iodoacetamide. The beads were then washed with mass-spec grade water and were resuspended in mass spec grade 50% MeCN/50% water. The samples were digested with Lys-C protease for 16 hours and with Solu-Trypsin for 1 hour (47 ^0^C), then for 4 hours (37 ^0^C). The supernatants were quenched with formic acid and the magnetic beads were removed. Samples were dried via vacuum centrifugation and stored at -80 ^0^C.

### Proteomics liquid chromatography-tandem mass spectrometry (LC-MS/MS) analysis

Samples were solubilized in 1% trifluoroacetic acid. An EASY nLC ultra-pressure liquid chromatography system was used to elute peptides onto a Fusion Tribrid mass spectrometer (Thermo Scientific). MS1 profiling was performed in a 375–1600 m/z range at a resolution of 70000. MS2 fragmentation was carried out on the top 15 ions by using a 1.6 m/z window and a normalized collision energy of 29 using higher energy collision-induced dissociation (HCD) with a dynamic exclusion of 15 s.

### Proteomics Data Analysis

All of mass spectra were analyzed with MaxQuant software version 1.6.10.43. MS/MS spectra were searched against the Homo Sapiens Uniprot protein sequence database based on version June 16th, 2021. Carbamidomethylation of cysteines was searched for as a fixed modification. Oxidation of methionines and acetylation of protein N-terminal as well as O-GlcNAc proteins termed as HexNac(ST) in MaxQuant software were searched against as variable modification. Enzyme was set to trypsin and LysC in a specific mode. All other parameters were used as default in MaxQuant. Label-free quantification was selected for group-specific parameters. Using Perseus, all contaminates identified by MaxQuant (streptavidin, reversed proteins, peptides with sequences <=2, etc.) are filtered out. Then the data was categorically grouped, peptides filtered for presence in at least 3 out of 4 positive replicates, analyzed by t-test analysis and plotted in a Volcano plot. Note: the serum-starved cyt-GlycoID (30 min of labeling) only had two successful proteomics runs, so for this condition positive hits were assigned when peptides were present in both replicates. Hits were scored by t-test analysis (p < 0.05) and fold-change (log2 > +/- 0.05). The full, processed data are presented in **Appendix Tables 1** and **2**. The mass spectrometry proteomics data have been deposited to the ProteomeXchange Consortium via the PRIDE partner repository with the dataset identifiers PXD033026, PXD033043, PXD033044, PXD033062, PXD033063, and PXD033066.

## Supporting Information

Appendix Table 1 (.xlsx)

Appendix Table 2 (.xlsx)

Additional and extended methods, generation of protein constructs, protein labeling optimization, proteomics sample preparation, insulin stimulation, analysis of non-sugar targeted labeling, analysis of serum-stimulated functional labeling, full STRING plots for all reported labeling experiments, supplemental discussion: contingency test for GlycoID-directed O-GlcNAc labeling. (PDF)

## Conflicts of Interest

The authors declare no competing financial interest.

## Acknowledgements

This study was supported by NIH R35GM142637-01, awarded to CF. The NIH:NCI Cancer Center Grant P30CA022453 to the Karmanos Cancer Institute supported contributions from the Wayne State Proteomics Core. We thank the Suzanne Walker Lab (Harvard Medical School) for the kind gift of OGT inhibitor OSMI-4. The plasmids 3xHA-miniTurbo-NLS_pCDNA3 (Addgene plasmid #107172; http://n2t.net/addgene:107172; RRID:Addgene_107172) and V5-miniTurbo-NES_pCDNA3 were gifts from Alice Ting (Addgene plasmid #107170; http://n2t.net/addgene:107170; RRID:Addgene_107170). We also thank Doug Haslitt for contributing preliminary bacterial data to the overall project.

## Notes

### Competing Interest Statement

The authors have declared no competing interest.

