## Supplementary Information for "Spatiotemporal proximity labeling tools to track GlcNAc sugar-modified functional protein hubs during cellular signaling"

Separate Excel files for proteomics data: **Appendix Tables 1 and 2**

**General:** The equipment, chemicals, and kits used in this study are shown in **Supplementary Table 1**.

*Supplementary Table 1:* ***General Items and Suppliers***

| **Instruments** | **Supplier** | **Catalogue Number** |
|---|---|---|
| MyCycler Thermal Cycler | BioRad | 170-9703 |
| NanoDrop-ND-1000 Spectrophotometer | ThermoFisher | |
| iBlot 2 dry blotting system | ThermoFisher | IB21001 |
| iBright$^{TM}$ FL1500 instrument | ThermoFisher | A44115 |
| **Consumables** | **Supplier** | **Catalogue Number** |
| T-75 flask | USAScientific | CC7682-4875 |
| 100 mm dish | FisherScientific | FB0875713 |
| 6-well dish | FisherScientific | 07-200-83 |
| 12-well dish | Corning | 3512 |
| NuPAGE™ 4 to 12%, Bis-Tris, 1.0–1.5 mm, Mini Protein Gels | ThermoFisher | NP0321BOX |
| iBlot™ 2 Transfer Stacks, nitrocellulose, mini | ThermoFisher | IB23002 |
| **Cells** | **Supplier** | **Catalogue Number** |
| XL10-Gold Ultracompetent cells | Agilent | 200314 |
| HeLa | ATCC | CRM-CCL-2 |
| HEK293T | ATCC | ACS-4500 |
| **Chemicals** | **Supplier** | **Catalogue Number** |
| V5-miniTurbo-NES_pCDNA3 | Addgene and Alice Ting | Addgene plasmid #107170; http://n2t.net/addgene:107170; RRID: Addgene_107170 |
| 3xHA-miniTurbo-NLS_pCDNA3 | Addgene and Alice Ting | Addgene plasmid #107172; http://n2t.net/addgene:107172; RRID: Addgene_107172 |
| Calf intestinal alkaline phosphatase (Quick CIP) | NEB | M0525 |
| T4 DNA Ligase | NEB | M0202 |
| DMEM | Sigma Aldrich | D6429 |
| HyClone Fetal Bovine Serum | Cytiva | SH30396.03 |
| HyClone Penicillin-Streptomycin solution | Cytiva | SV30010 |
| PBS pH 7.4 (1X) | ThermoFisher | 10010-023 |
| Trypsin 0.25% (1X) solution | Cytiva | SV30031.01 |
| TransIT-LT1 transfection reagent | Mirus Bio LLC | MIR 2304 |
| Bovine serum albumin | Research Products International | A30075 |
| Biotin | Carbosynth | 58-85-5 |
| Dharmacon$^{TM}$ ON-TARGET plus SMART pool human OGT siRNA | Horizon | L-019111-00-0005 |
| SMART pool human OGA siRNA | Horizon | L-012805-00-0005 |
| ON-TARGET control pool non-targeting pool siRNA | Horizon | D001810-10-05 |

| | | |
|---|---|---|
| Streptavidin-coated magnetic beads | NEB | S1410S |
| DTT (Dithiothreitol) | GoldBio | 27565-41-9 |
| iodoacetamide | Sigma | 16125 |
| MeCN (acetonitrile) | Fisher Chemical | 75-05-8 |
| Water | Fisher Chemical | 7732-18-5) |
| Lys-C protease | ThermoScientific | 90051 |
| SOLu-Trypsin | Sigma | EMS0004 |
| Formic acid | Thermo Scientific | 85178 |
| Formamide | OmniPur | 75-12-7 |
| SDS | FisherScientific | BP166-500 |
| **Kits** | **Supplier** | **Catalogue Number** |
| QIAquick PCR Purification Kit | Qiagen | 28104 |
| GeneJet Plasmid Miniprep Kit | ThermoFisher | K0502 |
| GeneJet Gel Extraction Kit | ThermoFisher | K0691 |
| Pierce Rapid Gold BCA Protein Assay Kit | ThermoFisher | A53225 |
| Pierce$^{TM}$ Quantitative Fluorometric Peptide Assay kit | ThermoFisher | 23290 |

**Generation of Constructs:** The full-length GafD gene was synthesized by ThermoFisher GeneArt from the reported template[1] (Uniprot: Q47341), as shown below. The O-GlcNAc binding domain (residues 23-178), which we termed "GafD_short," was specifically chosen for insertion using the primers listed in **Supplementary Table 2**. The constructs were created via sub-cloning our insert into vectors created by the Ting group:[2] V5-miniTurbo-NES_pCDNA3 (Addgene plasmid #107170) and 3xHA-miniTurbo-NLS_pCDNA3 (Addgene plasmid #107172). The target GafD gene was generated through amplification via PCR using MyCycler Thermal Cycler (BioRad) and purified using 1.5% agarose gel (100 V for 40 minutes). The amplified PCR products were purified via QIAquick PCR Purification Kit (Qiagen, 28104). Plasmids were isolated using GeneJet Plasmid Miniprep Kit (ThermoFisher, K0502), with the use of a NanoDrop ND-1000 Spectrometer (ThermoFisher) to confirm their concentrations. The miniTurbo containing plasmids and PCR products were double digested with the restriction enzymes (double check, NEB) and the plasmid was dephosphorylated using calf intestinal alkaline phosphatase (Quick CIP, NEB: M0525); both were purified using a 0.8% agarose gel and cleaned up via GeneJet Gel Extraction Kit (ThermoFisher, K0691). The plasmid containing miniTurboID and the GafD gene were ligated using T4 DNA Ligase (NEB, M0202) at 16 $^0$C, overnight with shaking (700 rpm). Plasmids were transformed into XL10-Gold Ultracompetent cells (Agilent, 200314). Confirmation of gene insertion was completed using PCR. Cloning was verified by Sanger DNA sequencing. The sequences generated in this study are collected in **Supplemental Table 3**. Addgene ID's 184640 (cyt-GlycoID) and 184641 (nuc-GlycoID).

*Supplementary Table 2: Primers used in this study*

| **Primer Name** | **Sequence** |
|---|---|
| HindIII_GafD_short_for | **CCCCCC AAGCTT** ATG GCC GTG TCC TTC ATC GG |
| HindIII_GafD_short_rev | **CCCCCC AAGCTT** CAG AGC TGC CAT TGT AGG G |
| NotI_GafD_short_rev | **CCCCCC GCGGCCGC** AGA GCT GCC ATT GTA GGG |

GeneArt construct:

```
     M  T  N  F  Y  K  V  C  L  A  V  F  I  L  V  C  C  N  I  S  H  A  A
1.   ATGACCAACTTCTATAAAGTTTGCCTGGCCGTTTTTATTCTGGTGTGTTGTAATATTAGCCATGCAGCC
     V  S  F  I  G  S  T  E  N  D  V  G  P  S  Q  G  S  Y  S  S  T  H  A
70.  GTTAGCTTTATTGGTAGCACCGAAAATGATGTTGGTCCGAGCCAGGGTAGCTATAGCAGCACCCATGCA
     M  D  N  L  P  F  V  Y  N  T  G  Y  N  I  G  Y  Q  N  A  N  V  W  R
139. ATGGATAATCTGCCGTTTGTGTATAACACCGGCTATAACATTGGTTATCAGAATGCAAATGTGTGGCGT
     I  S  G  G  F  C  V  G  L  D  G  K  V  D  L  P  V  V  G  S  L  D  G
208. ATTAGCGGTGGTTTTTGTGTTGGTCTGGATGGTAAAGTTGATCTGCCGGTTGTTGGTAGCCTGGATGGT
     Q  S  I  Y  G  L  T  E  E  V  G  L  L  I  W  M  G  D  T  N  Y  S  R
277. CAGAGCATTTATGGTCTGACCGAAGAAGTGGGTCTGCTGATTTGGATGGGTGATACCAATTATAGCCGT
     G  T  A  M  S  G  N  S  W  E  N  V  F  S  G  W  C  V  G  N  Y  V  S
346. GGCACCGCAATGAGCGGTAATAGCTGGGAAAATGTTTTTAGCGGTTGGTGCGTTGGTAATTATGTTAGC
     T  Q  G  L  S  V  H  V  R  P  V  I  L  K  R  N  S  S  A  Q  Y  S  V
415. ACCCAGGGTCTGAGCGTTCATGTTCGTCCGGTTATTCTGAAACGTAATAGCAGCGCACAGTATAGCGTT
     Q  K  T  S  I  G  S  I  R  M  R  P  Y  N  G  S  S  A  G  S  V  Q  T
484. CAGAAAACCAGCATTGGTAGTATTCGTATGCGTCCGTATAATGGTAGCAGTGCAGGTAGCGTGCAGACC
     T  V  N  F  S  L  N  P  F  T  L  N  D  T  V  T  S  C  R  L  L  T  P
553. ACCGTGAATTTTAGCCTGAATCCGTTTACACTGAATGATACCGTTACCAGCTGTCGTCTGCTGACCCCG
     S  A  V  N  V  S  L  A  A  I  S  A  G  Q  L  P  S  S  G  D  E  V  V
622. AGCGCAGTTAATGTTAGCCTGGCAGCAATTAGCGCAGGTCAGCTGCCGAGCAGCGGTGATGAAGTTGTT
     A  G  T  T  S  L  K  L  Q  C  D  A  G  V  T  V  W  A  T  L  T  D  A
691. GCAGGTACAACCAGCCTGAAACTGCAGTGTGATGCGGGTGTTACCGTTTGGGCAACCCTGACCGATGCA
     T  T  P  S  N  R  S  D  I  L  T  L  T  G  A  S  T  A  T  G  V  G  L
760. ACCACACCGAGCAATCGTAGCGATATTCTGACCCTGACAGGTGCAAGCACCGCAACAGGTGTTGGCCTG
     R  I  Y  K  N  T  D  S  T  P  L  K  F  G  P  D  S  P  V  K  G  N  E
829. CGTATTTACAAAAATACCGATAGCACACCGCTGAAATTTGGTCCGGATAGTCCGGTTAAAGGTAATGAA
     N  Q  W  Q  L  S  T  G  T  E  T  S  P  S  V  R  L  Y  V  K  Y  V  N
898. AATCAGTGGCAGCTGAGCACCGGCACCGAAACCAGTCCGAGCGTTCGTCTGTATGTTAAATATGTGAAT
     T  G  E  G  I  N  P  G  T  V  N  G  I  S  T  F  T  F  S  Y  Q
967. ACAGGCGAAGGCATTAATCCGGGTACAGTTAATGGTATTAGCACCTTTACCTTCAGCTACCAG
```

*Supplementary Table 3: Construct Design Amino Acid Sequences*

| | |
|---|---|
| V5 tag | GKPIPNPLLGLDST |
| HA tag | YPYDVPDYA |
| GafD-short | MAVSFIGSTENDVGPSQGSYSSTHAMDNLPFVYNTGYNIGYQNAN VWRISGGFCVGLDGKVDLPVVGSLDGQSIYGLTEEVGLLIWMGDT NYSRGTAMSGNSWENVFSGWCVGNYVSTQGLSVHVRPVILKRNS SAQYSVQKTSIGSIRMRPYNGSS |
| miniTurboID | MIPLLNAKQILGQLDGGSVAVLPVVDSTNQYLLDRIGELKSGDACIA EYQQAGRGSRGRKWFSPFGANLYLSMFWRLKRGPAAIGLGPVIGI VMAEALRKLGADKVRVKWPNDLYLQDRKLAGILVELAGITGDAAQI VIGAGINVAMRRVEESVVNQGWITLQEAGINLDRNTLAAMLIRELRA ALELFEQEGLAPYLSRWEKLDNFINRPVKLIIGDKEIFGISRGIDKQG ALLLEQDGVIKPWMGGEISLRSAEK |
| cyt-GafD-mTurboID-V5 (cyt-GlycoID) | MAVSFIGSTENDVGPSQGSYSSTHAMDNLPFVYNTGYNIGYQNAN VWRISGGFCVGLDGKVDLPVVGSLDGQSIYGLTEEVGLLIWMGDT NYSRGTAMSGNSWENVFSGWCVGNYVSTQGLSVHVRPVILKRNS SAQYSVQKTSIGSIRMRPYNGSSAAATMGKPIPNPLLGLDSTASIPL LNAKQILGQLDGGSVAVLPVVDSTNQYLLDRIGELKSGDACIAEYQ QAGRGSRGRKWFSPFGANLYLSMFWRLKRGPAAIGLGPVIGIVMA EALRKLGADKVRVKWPNDLYLQDRKLAGILVELAGITGDAAQIVIGA GINVAMRRVEESVVNQGWITLQEAGINLDRNTLAAMLIRELRAALE LFEQEGLAPYLSRWEKLDNFINRPVKLIIGDKEIFGISRGIDKQGALL LEQDGVIKPWMGGEISLRSAEKLPPLERLTL |
| nuc-GafD-mTurboID-HA (nuc-GlycoID) | MAVSFIGSTENDVGPSQGSYSSTHAMDNLPFVYNTGYNIGYQNAN VWRISGGFCVGLDGKVDLPVVGSLDGQSIYGLTEEVGLLIWMGDT NYSRGTAMSGNSWENVFSGWCVGNYVSTQGLSVHVRPVILKRNS SAQYSVQKTSIGSIRMRPYNGSSAAATMYPYDVPDYAGYPYDVPD YAGYPYDVPDYAASIPLLNAKQILGQLDGGSVAVLPVVDSTNQYLL DRIGELKSGDACIAEYQQAGRGSRGRKWFSPFGANLYLSMFWRL KRGPAAIGLGPVIGIVMAEALRKLGADKVRVKWPNDLYLQDRKLAG ILVELAGITGDAAQIVIGAGINVAMRRVEESVVNQGWITLQEAGINLD RNTLAAMLIRELRAALELFEQEGLAPYLSRWEKLDNFINRPVKLIIG DKEIFGISRGIDKQGALLLEQDGVIKPWMGGEISLRSAEKPKKKRKV DPKKKRKVDPKKKRKV |
| NES tag | LPPLERLTL |
| NLS tag | PKKKRKV |

**Mammalian Cell Culture and Transfection**: Cells were obtained from ATCC. All consumables (pipette tips, glass Pasteur pipettes, Eppendorf tubes) were sterilized via autoclave. HeLa cells were cultured in DMEM (Sigma Aldrich, D6429) supplemented with

10% (v/v) HyClone Fetal Bovine Serum (Cytiva, SH30396.03) and 1% HyClone Penicillin-Streptomycin solution (Cytiva, SV30010) at 37 $^0$C under 5% $CO_2$. All mammalian cell manipulations were done inside a laminar flow hood that was sterilized with 70% ethanol. To seed cells, the cells were carefully washed with sterile 7 mL of PBS pH 7.4 (1X) (ThermoFisher, 10010-023). The 1.5 mL of Trypsin 0.25% (1X) solution (Cytiva, SV30031.01) was added to the flask and incubated for 5 minutes at 37 $^0$C under 5% $CO_2$. The trypsin was then neutralized with serum containing growth media, where the trypsin can be further removed via centrifugation (300 x g for 3 minutes) in a sterile 15 mL centrifuge tube (FisherScientific, 14-955-237). The cell pellet was washed with PBS and recentrifuged. The cells can then be seeded into desired flask (2.1x$10^6$ cells for T-75 flask [USAScientific, CC7682-4875], 2.2x$10^6$ cells for 100 mm dish [FisherScientific, FB0875713], 0.3x$10^6$ cells for 6-well dish [FisherScientific, 07-200-83], 0.1 x$10^6$ cells for 12-well dish [Corning, 3512]). For transient transfection, cells were transfected using TransIT-LT1 transfection reagent (Mirus Bio LLC, MIR 2304) according to the manufacturer's protocol. The transfected cells were incubated for 48-72 hours before use. General reagents used in this study are collected in **Supplemental Table 4**.

*Supplementary Table 4: Transfection amounts*

| Culture Vessel | 24-well | 12-well | 6-well | 100 mm |
|---|---|---|---|---|
| Surface Area | 1.9 $cm^2$ | 3.8 $cm^2$ | 9.6 $cm^2$ | 59 $cm^2$ |
| Complete Growth Media | 0.5 mL | 1.0 mL | 2.5 mL | 15.5 mL |
| Serum-free Media | 50 µL | 100 µL | 250 µL | 1.5 mL |
| DNA (1 µg/µL) | 0.5 µL | 1 µL | 2.5 µL | 15 µL |
| *Trans*IT-LT1 Reagent | 1.5 µL | 3 µL | 7.5 µL | 45 µL |

**Western Blot:** To analyze cell lysates via immunoblot, cells were collected with a cell scraper from plates in RIPA buffer containing protease inhibitors (150 mM NaCl, 1% Nonidet P-40, 0.5% Na-deoxycholate, 0.1% sodium dodecyl sulfate [SDS], and 50 mM Tris-pH 7.4). Cell lysates were briefly sonicated and centrifuged (12,000 x g for 10 minutes at 4 $^0$C) to collect the soluble protein fraction. Protein concentration was determined via Pierce Rapid Gold BCA Protein Assay Kit (ThermoFisher, A53225). Samples were boiled in SDS gel-loading buffer for 5 minutes. Proteins were separated on a 4-12% gradient gel (NuPAGE™ 4 to 12%, Bis-Tris, 1.0–1.5 mm, Mini Protein Gels; ThermoFisher, NP0321BOX) and transferred to a nitrocellulose membrane (iBlot™ 2 Transfer Stacks, nitrocellulose, mini; ThermoFisher, IB23002) using an iBlot 2 dry blotting system (ThermoFisher, IB21001). After blocking with 5% w/v bovine serum albumin (Research Products International, A30075) in TBST buffer (10 mM Tris-pH8, 150 mM NaCl, 0.05% Tween 20) for 1 hour, the membrane was incubated with the appropriate antibody following the manufacture's protocol. The signals from the antibodies were detected via iBright™ FL1500 instrument (ThermoFisher, A44115). To detect biotinylated proteins, the membranes were incubated with Cy5- or HRP-conjugated streptavidin. All antibodies used in this study are collected in **Supplemental Table 5**.

*Supplementary Table 5: Antibodies used in this study*

| Target | Conjugate | Host | Supplier | Dilution |
|---|---|---|---|---|
| Biotin | HRP | Goat | CST – 7075S | 1:1000-1:3000 |
| Biotin | Cy5 | Bacterial | Invitrogen – SA1011 | 1:1000 |
| Flag | N/A | Mouse | Sigma – F3165 | 1:1000 |
| HA | N/A | Mouse | BioLegend – 901503 | 1:1000 |
| HA | N/A | Rabbit | CST – 3724S | 1:1000 |
| Mouse IgG | HRP | Goat | Thermo – A28175 | 1:10000-1:200000 |
| OGT | N/A | Rabbit | CST – 24083S | 1:1000 |
| Rabbit IgG | HRP | Donkey | Thermo – 31458 | 1:10000-1:200000 |
| V5 | N/A | Rabbit | CST – 13202S | 1:1000 |
| V5 | N/A | Rabbit | Thermo – PA1-993 | 1:1000-1:5000 |
| Rabbit IgG | Alexa Fluor 555 | Goat | **A27039** | 1:1000 |
| Rabbit IgG | Alexa Fluor 488 | Goat | **A-11008** | 1:1000 |
| O-GlcNAc MultiMab | N/A | Rabbit | CST – 82332S | 1:1000 |

**Biotin Labeling with Fusion Constructs**: For biotin labeling experiments of transiently transfected cells, biotin was added 48-72 hours after transfection. Biotin (Carbosynth, 58-85-5) was diluted to 100 μM (or desired concentration) in complete growth media and added directly to cells. The cells were incubated at 37 $^0$C for the desired amount of time. For both western blots and proteomics, labeling was stopped by washing with cold PBS and freezing at -80 $^0$C.

**Thiamet-G or OSMI Inhibition:** After plating cells and transfecting them with the appropriate vector and 16-24 hours before biotin labeling, the plates are incubated with inhibitor diluted into complete media (40 μM for OSMI, and 10 μM for Thiamet-G) overnight. Labeling was stopped with washing with PBS and freezing at -80 $^0$C.

**siRNA Knockdown:** For OGT knockdown, cells were transfected with Dharmacon™ ON-TARGET plus SMART pool human OGT siRNA (# L-019111-00-0005), SMART pool human OGA siRNA (# L-012805-00-0005), or ON-TARGET control pool non-targeting pool siRNA (#D001810-10-05) as control using DharmaFECT™ transfection reagent, as described by the manufacturer. The SMART pool consists of a combination of 4 different siRNA oligomers optimized for knockdown in human cell lines, which were used here instead of two distinct siRNA sequences for OGT knockdown and its corresponding SMART pool control knockdown. The dried siRNA pellets were recentrifuged and resuspended in RNase-free 1x siRNA buffer (60 mM KCl, 6 mM HEPES-pH 7.5, and 0.2 mM $MgCl_2$) to final concentration of 20 μM and aliquoted into 20 μL samples. Plates were seeded to the desired confluency. For transfection, the siRNA aliquot was diluted to 5 μM and transfected using the amounts found in **Supplemental Table 6**.

Supplementary Table 6: RNA knockdown reagents used in this study

| | | Tube 1: diluted siRNA | | Tube 2: diluted DharmaFECT | | | |
|---|---|---|---|---|---|---|---|
| **Plating Format** | Surface Area ($cm^2$/well) | Volume of 5 µM siRNA (µL) | Serum-free media (µL) | Volume of DharmaFECT reagent (µL) | Serum-free media (µL) | Complete media (µL/well) | Total transfection volume (µL/well) |
| **96** | 0.3 | 0.5 | 9.5 | 0.05-0.5 | 9.95-9.5 | 80 | 100 |
| **24** | 2 | 2.5 | 47.5 | 0.25-2.5 | 49.75-47.5 | 400 | 500 |
| **12** | 4 | 5 | 95 | 0.5-5.0 | 99.5-95.0 | 800 | 1000 |
| **6** | 10 | 10 | 190 | 1.0-10.0 | 199.0-190.0 | 1600 | 2000 |

For 24-well plates, 2 µL of DharmaFECT reagent was sufficient for this siRNA transfection. The reagents were gently mixed via pipetting and incubated for 5 minutes at room temperature. The tubes were then combined and incubated for an additional 20 minutes. The media was removed from the plate and replaced with the appropriate amount of transfection reagent. The transfected plates were incubated at 37 $^0$C under 5% $CO_2$ for 48-96 hours for protein analysis.

**Sample Preparation for Proteomics**: Cells were cultured in d-100 mm TC-treated petri dishes. All cells were transiently expressing desired construct. All cells were labeled with 100 µM biotin using aforementioned methods. Labeling was stopped by washing with cold PBS and freezing at -80 $^0$C. The cells were detached from the plate via scrapper with lysis buffer (150 mM NaCl, 0.5 mM tris, 1% NP40, 0.1% SDS) and collected in Eppendorf tubes. The cells were lysed via passage through needle (at least 10 passes) or sonication and clarified with centrifugation at 10,000 x g for 10 minutes at 4 $^0$C.

For enrichment of biotinylated proteins, 100 µL of streptavidin-coated magnetic beads (NEB S1410S) were washed twice with RIPA buffer and then incubated with clarified lysates (400 µg protein) with rotation at 4 $^0$C overnight. The magnetic beads were then washed once with 500 µL RIPA buffer, once with 500 µL wash buffer (50 mM Tris, ph 7.4, 2% SDS), and twice with 500 µL RIPA buffer. Magnetic beads were resuspended in 500 μL 10 mM DTT (Dithiothreitol, GoldBio 27565-41-9) in PBS at 37 $^0$C for 30 minutes, which was then cooled to room temperature. The supernatant was discarded. The magnetic beads were then resuspended in 1 mL 30 mM iodoacetamide (Sigma, 16125) (protect from light) at r.t.  for 30 minutes. The supernatant was discarded, and the beads were washed with pure mass-spec grade water. The magnetic beads were resuspended in 300 μL 50% MeCN/50% water (Fisher Chemical, 75-05-8; Fisher Chemical, 7732-18-5) (ms-grade). The proteins were then digested with Lys-C protease (Thermo Scientific, 90051) with a 1:100 ratio Lys-C to protein sample (~0.3 μL for 50 μL resin) at 37 $^0$C for 16 hours without shaking. The proteins were further digested with SOLu-Trypsin (Sigma, EMS0004) at a ratio 1:20 trypsin weight to sample weight (50 μL resin, 3 μL Trypsin) at 47 $^0$C for one hour, then cooled to 37 $^0$C for four hours with rotation. The digestion was quenched by bringing mixture to a final concentration of 1% formic acid (Thermo Scientific, 85178). The beads were removed from the mixture via magnet (or centrifugation) and washed the beads twice with 200 μL 50% MeCN and once M.S.-water. Bead fragments were removed with centrifugation (10,000 x g for 10 minutes). Samples were concentrated via speedvac set to 40 $^0$C and the residues were stored at -80 $^0$C. Peptide concentrations were determined via Pierce$^{TM}$ Quantitative Fluorometric Peptide Assay kit (ThermoFisher, 23290), following the manufacturer's protocol. The detection of detergents was conducted using a SDS assay, suing Stains-all dye. A stock solution of 1.8 mM stains-all was made using 50% propanol:water (protect from light) (e.g. 10 mL solution needs 10 mg of stains-all). A 90 μM working solution was diluted from the stock solution in 5% formamide (OmniPur, 75-12-7) (e.g. for 5 mL; mix 0.25 mL stock, 0.25 mL formamide, 4.5 mL water, 2.5% propanol final). This solution can be stored at room temperature in the dark for ~four days. Pipette 1 μL of sample and 1 μL of a standard curve SDS (FisherScientific, BP166-500) sample (0.02-0.1%) into a 96 well plate. Standard curve used started at 0.02% with increments of 0.01% (e.g. 0.02, 0.03, 0.04, etc.). 200 μL of the working solution was added into each well with a sample or standard (keep the plate protected from the light). The plate was read using a plate reader at 445 nm. The samples should have a minimum amount of SDS to prevent damage to the mass spectrum column.

**Proteomics**: All of mass spectra were analyzed with MaxQuant software version 1.6.10.43. MS/MS spectra were searched against the Homo Sapiens Uniprot protein sequence database based on version June 16th, 2021. Carbamidomethylation of cysteines was searched for as a fixed modification. Oxidation of methionines and acetylation of protein N-terminal as well as O-GlcNAc proteins termed as HexNac(ST) in MaxQuant software were searched against as variable modification. Enzyme was set to trypsin and Lys-C in a specific mode. All other parameters were used as default in MaxQuant. Label-free quantification was selected for group-specific parameters. Using Perseus, all contaminates identified by MaxQuant (streptavidin, reversed proteins, peptides with sequences <=2, etc.) are filtered out. Then the data is categorically grouped, using data with at least 3 out of 4 positive replicates to do a T-Test and plot in a Volcano plot. Note: the serum-starved cyt-GlycoID (30 min of labeling) only had two successful proteomics runs, so its positive hits were 2 out of 2 replicates.

**Data Analysis**

The protein lists for each labeling condition were collected in Supplementary Table 1 and Supplementary Table 2, attached as separate .xlsx files. For each condition, protein hits that were exclusive to each condition were combined with the significantly enriched proteins identified by Volcano plot analysis. These total lists for each condition were cross-referenced with the dataset from the O-GlcNAcome website (https://www.oglcnac.mcw.edu/), as published.[3] Proteins were also analyzed against the OGT Protein Interaction Network (OGT-PIN) downloaded from the OGT-PIN website (https://oglcnac.org/ogt-pin/), as published.[4] Total interactome analysis was performed using the STRING online protein-protein association network database (https://string-db.org/ version 11.5) with the following parameter settings:

**Input**

Multiple proteins

Organism: Homo sapiens

---

**Settings**

**Network type**: Physical Subnetwork

**Meaning of network edges**: evidence

**Active interaction sources**: Experiments only (“textmining” and “databases” were de-selected)

**Minimum required interaction score**: medium confidence (0.400)

Max number of interactors to show: 1st shell: query proteins only / 2nd shell: none

---

**Clusters**

**Clustering options**: kmeans clustering

**Number of clusters**: varied until significance was not met; 0-6 clusters were identified for each condition.

**Significance**: $p \leq 0.05$

**Supplemental Discussion: Contingency test for GlycoID-directed O-GlcNAc labeling**

A contingency test was used to estimate the probability that the GlycoID tools labeled O-GlcNAcylated proteins more frequently than random proteins. The labeling results were split into two categories, known O-GlcNAc proteins vs. proteins without known O-GlcNAc modifications. The null hypothesis was that GlycoID does not label O-GlcNAc frequencies at greater levels than any protein. A Fisher’s exact test was used due to relatively small sample sizes. 2x2 continency tables were used to test whether if it was possible to reject this null hypothesis. The test and result are shown in **Supplemental Table 7**.

*Supplementary Table 7: Contingency Table for GlycoID-based labeling of known O-GlcNAc proteins*

| | nuc-mTurbo | nuc-GlycoID |
|---|---|---|
| O-GlcNAc protein | 17 | 21 |
| Non-GlcNAcylated | 35 | 48 |

Fisher’s exact test (1-tailed) p = 0.41 (NS)

| | cyt-mTurbo | cyt-GlycoID |
|---|---|---|
| O-GlcNAc protein | 5 | 50 |
| Non-GlcNAcylated | 9 | 52 |

Fisher’s exact test (1-tailed) p = 0.259 (NS)

However, since proximity labeling experiments also label proteins that physically associate with the target, analysis was extended to include the known GlcNAc interactome. Fisher's exact test was applied to the known O-GlcNAc protein interactome for non-targeted mTurbo labeling GlycoID vs. GlycoID enrichments results. The resulting categories were split between Group 1 + 2 proteins (O-GlcNAcylated proteins with their known physical interactomes) and Group 3 + 4 proteins (non-O-GlcNAc interactomes). The test and result are shown in **Supplemental Table 8**.

*Supplementary Table 8: Contingency Table for GlycoID-based labeling of known O-GlcNAc proteins + interactomes*

| | nuc-mTurbo | nuc-GlycoID |
|---|---|---|
| Group 1 + 2 | 26 | 49 |
| Group 3 + 4 | 24 | 20 |

Fisher's exact test (1-tailed) p = 0.0340

| | cyt-mTurbo | cyt-GlycoID |
|---|---|---|
| Group 1 + 2 | 5 | 81 |
| Group 3 + 4 | 9 | 21 |

Fisher's exact test (1-tailed) p = 0.00140

The results suggest that GlycoID proximity labeling strategy effectively labeled O-GlcNAc-modified protein clusters in HeLa cells. Care had to be taken interpret results that included the known protein-protein interactions of O-GlcNAc proteins. Otherwise, non-O-GlcNAc interaction partners confounded contingency test analyses based strictly on "known O-GlcNAc modification" state as a categorical variable.

**Supporting Figures:**

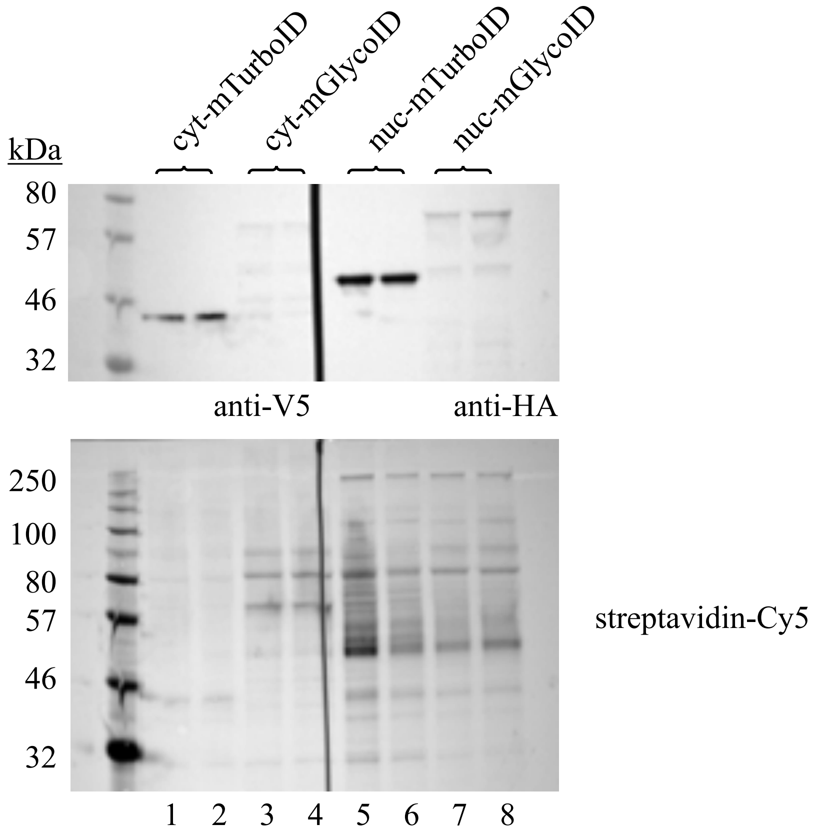


**Supplemental Figure 1:** Western blot for the expression of cyt-mTurboID (1, 2), cyt-GlycoID (3, 4), nuc-mTurboID (5, 6), nuc-GlycoID (7, 8) in HEK293T cells. All experiments used 100 μM biotin and were allowed to label for 6 hours.

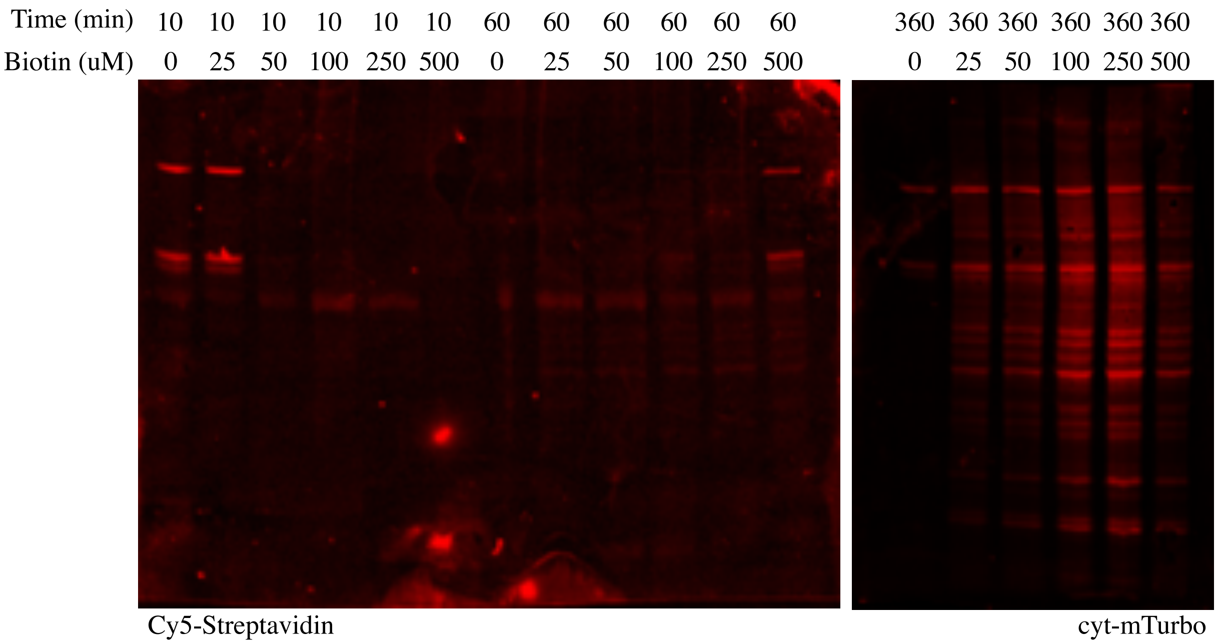


**Supplemental Figure 2A**: Western blots (Cy5-Streptavidin, 1:1000) for the biotin (0-500 µM) and time (10 min. to 6 hours) screen for cyt-mTurbo in HeLa cells. Blots were imaged using iBright™ FL1500 instrument.

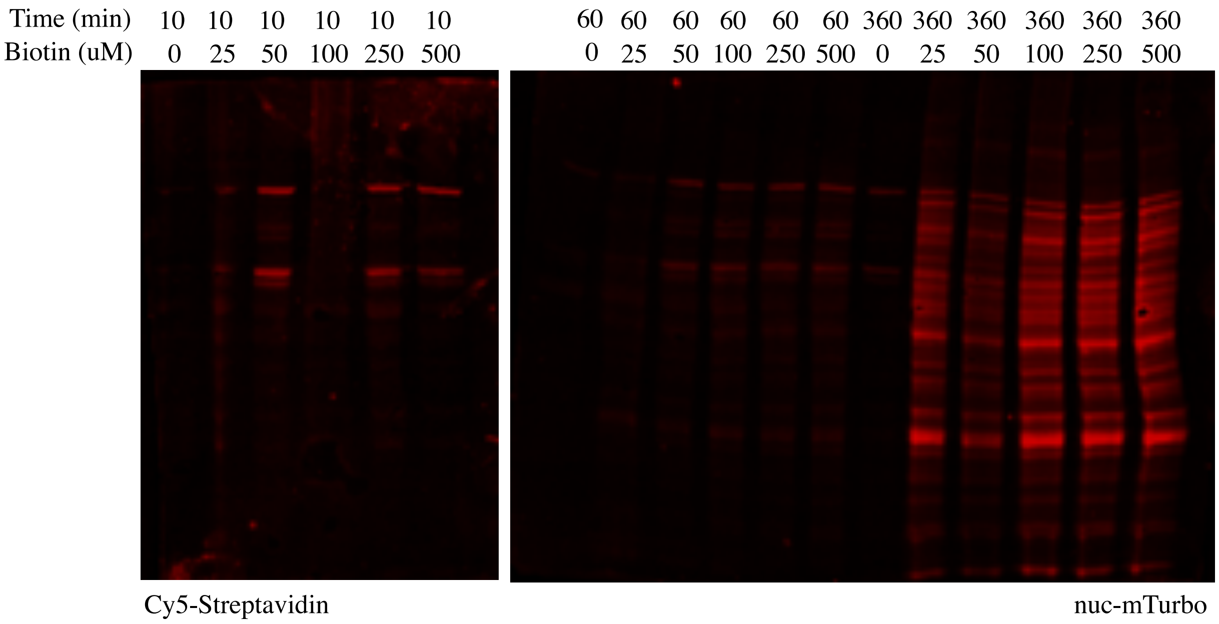


**Supplemental Figure 2B**: Western blots (Cy5-Streptavidin, 1:1000) for the biotin (0-500 µM) and time (10 min. to 6 hours) screen for nuc-mTurbo in HeLa cells. Blots were imaged using iBright™ FL1500 instrument.

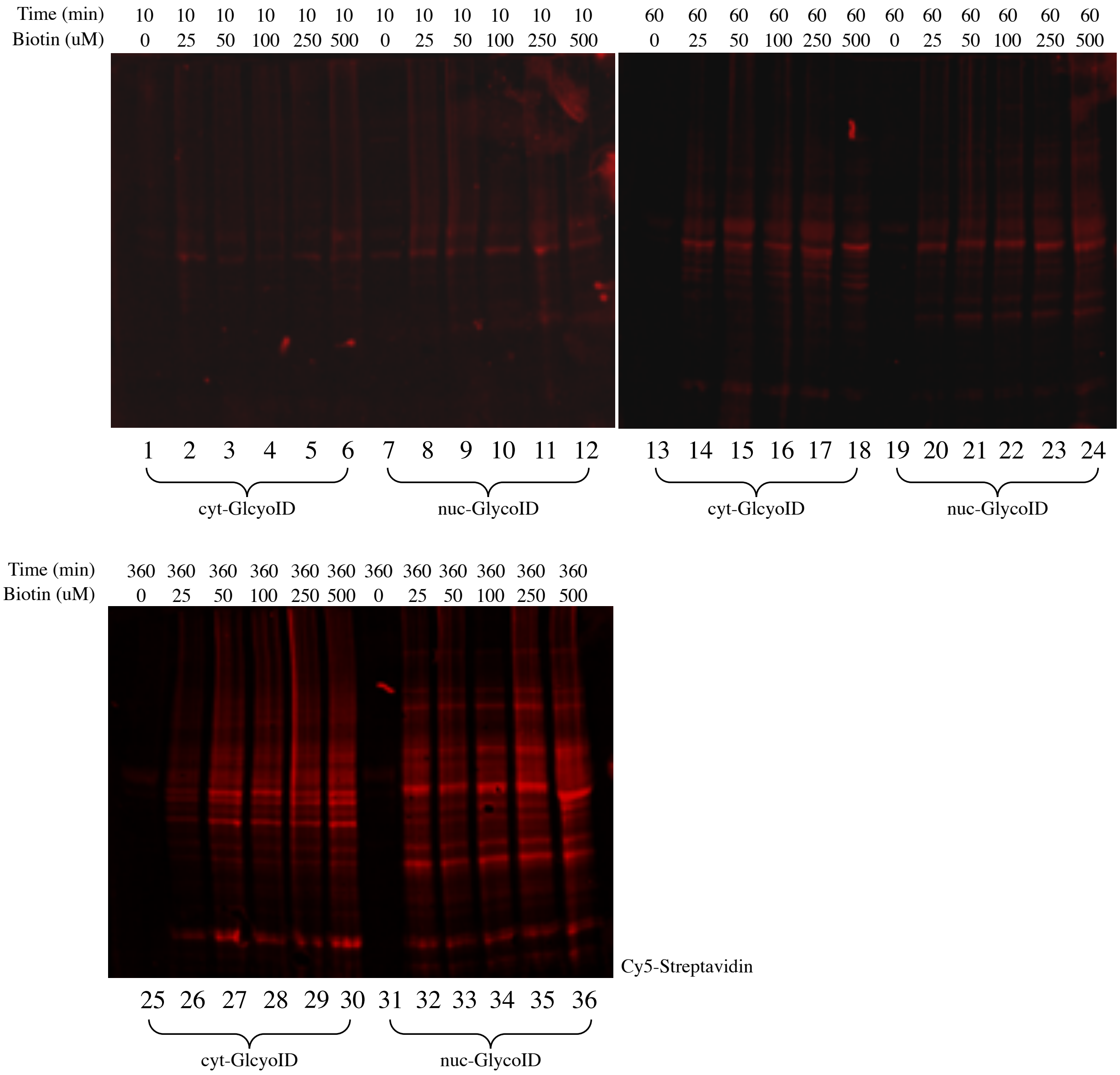


**Supplemental Figure 2C**: Western blots (Cy5-Streptavidin, 1:1000) for the biotin and time screen for cyt-GlycoID (1-6, 13-18, 25-30) or nuc-GlycoID (7-12, 19-24, 31-36) in HeLa cells. Blots were imaged using iBright™ FL1500 instrument.

a)

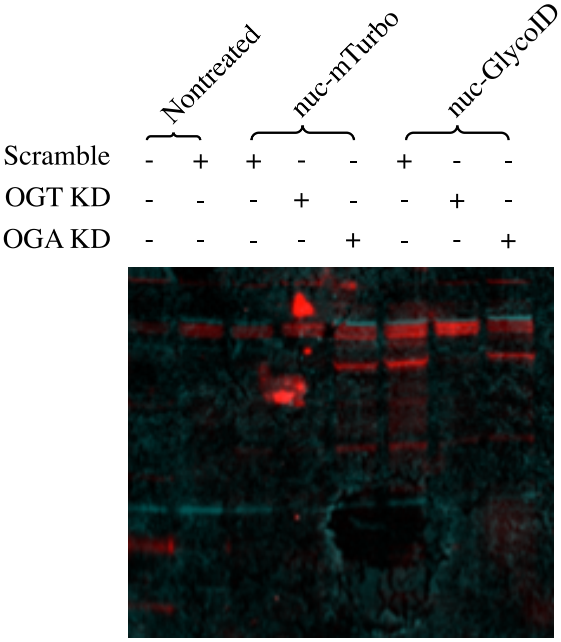


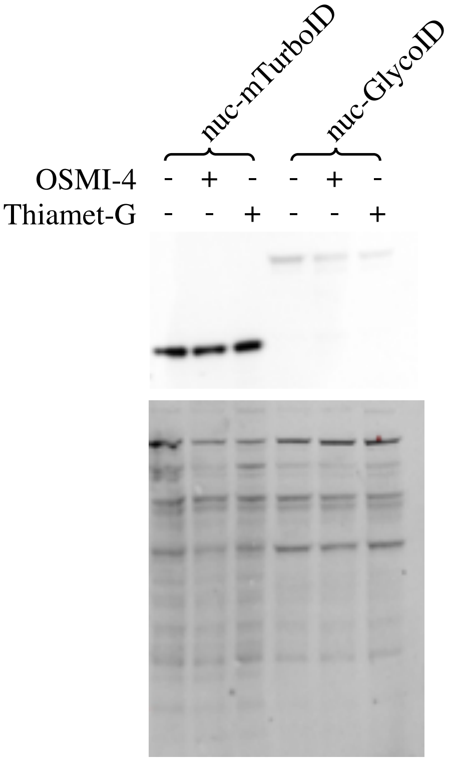


c)

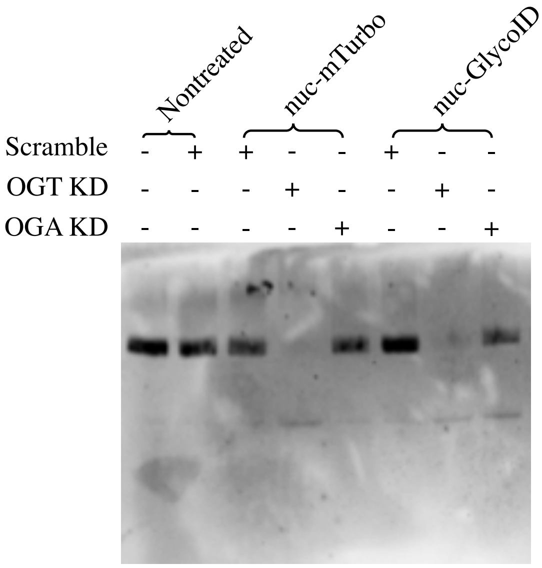


b)

**Supplemental Figure 3:** Whole blots for inhibition studies using GlycoID. a) Western blot for the activity of nuc-Turbo, and nuc-GlycoID blotted against O-GlcNAc MultiMab (1:1000)/Anti-Rabbit-AlexaFluor-488 (Green, 1:1000) and Streptavidin-Cy5 (Red, 1:1000). b) Western blot of the knockdown of OGT/OGA using siRNA blotted against anti-OGT (1:1000)/Anti-Rabbit-HRP (1:10,000). c) Western blot for the inhibition of OGT/OGA using Thiamet-G (10 μM) and OSMI-4 (40 μM) blotted against anti-HA (1:1000)/Anti-Rabbit (1:10,000). All blots used 500 μM biotin with 6 hours allowed for labeling. Blots were imaged using Blots were imaged using iBright™ FL1500 instrument.

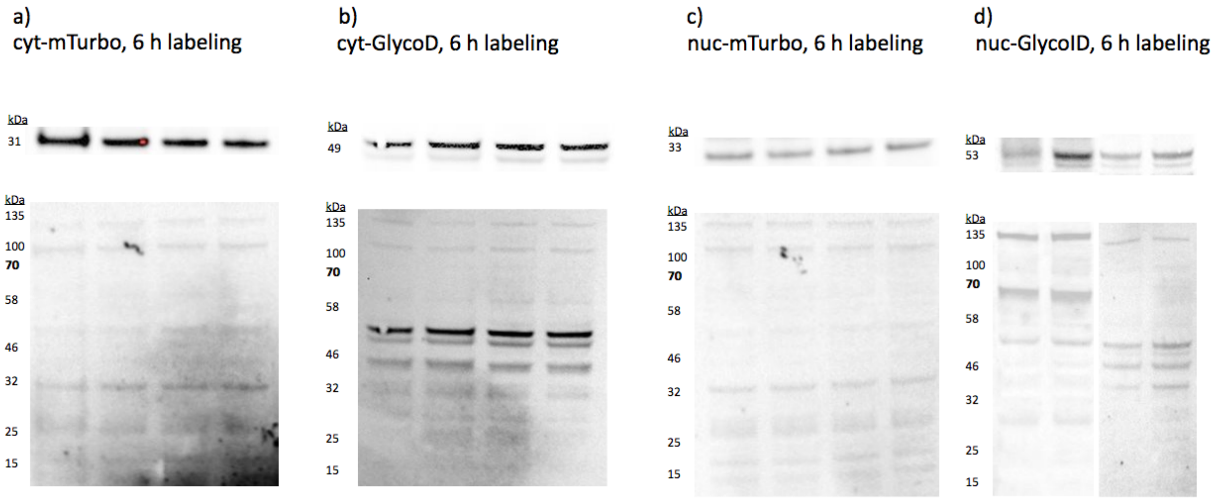


**Supplemental Figure 4:** Western blot for the expression of (a) cyt-mTurbo, (b) cyt-GlycoID, (c) nuc-mTurbo, and (d) nuc-GlycoID in HeLa cells used in proteomics experiments. Labeling was conducted for 6 hours with 100 μM biotin. Blots were imaged using iBright™ FL1500 instrument.

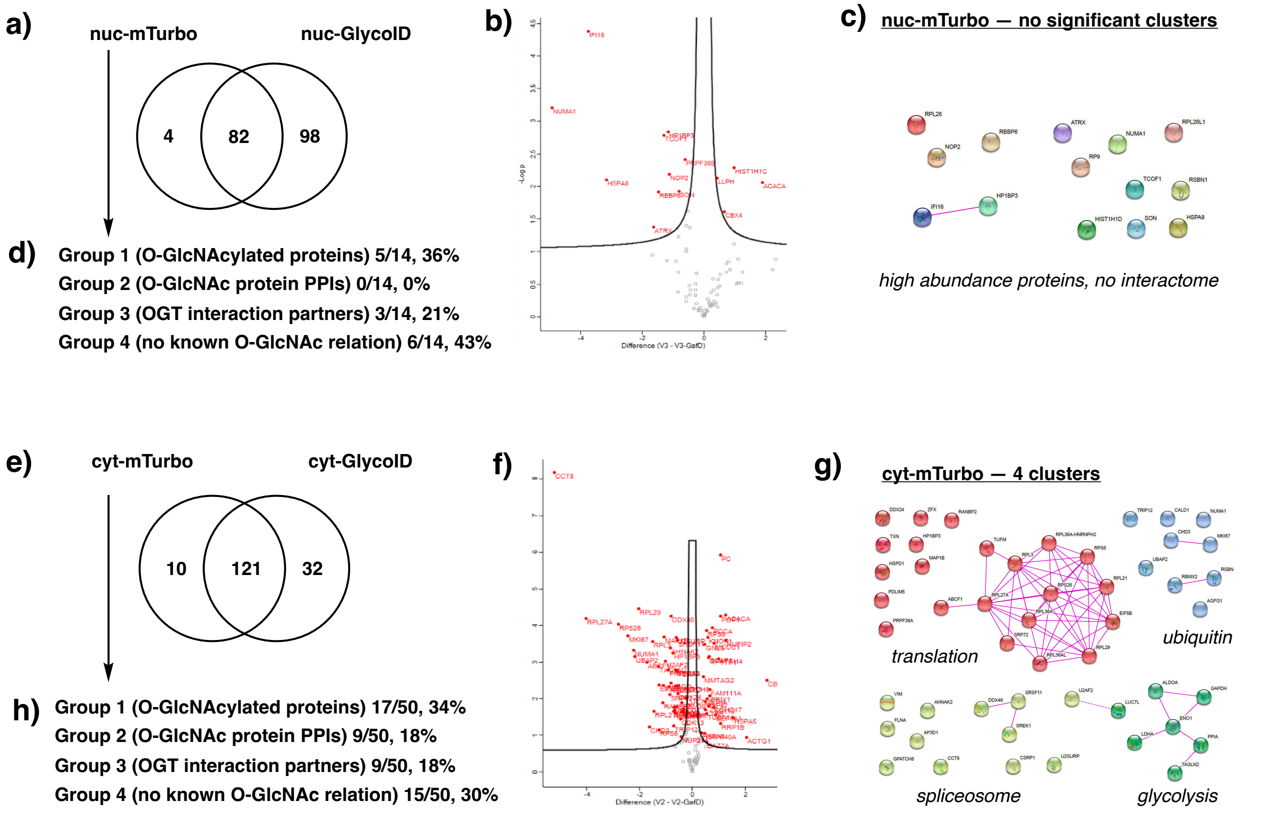


**Supplemental Figure 5 –** Analysis of nuc-mTurbo and cyt-mTurbo (non-sugar targeted) quantitative proteomics with GlycoID constructs. ***a***) Exclusive hits between non-targeted nuc-mTurbo vs. nuc-GlycoID. ***b***) Enrichment analysis between nuc-mTurbo and nuc-GlycoID, statistically significant hits are shown above the volcano plot cutoffs. ***c***) Physical interactions between nuc-GlycoID hits reveal functional clusters with key O-GlcNAc linkages. ***d***) The protein groups labeled by nuc-GlycoID, as defined in the main text. ***e***)-***h***) *A*nalysis for cyt-GlycoID vs. cyt-mTurbo. Full-sized STRING plots with labeled O-GlcNAc hits are found in **Supporting Figures 5** and **7-9**.

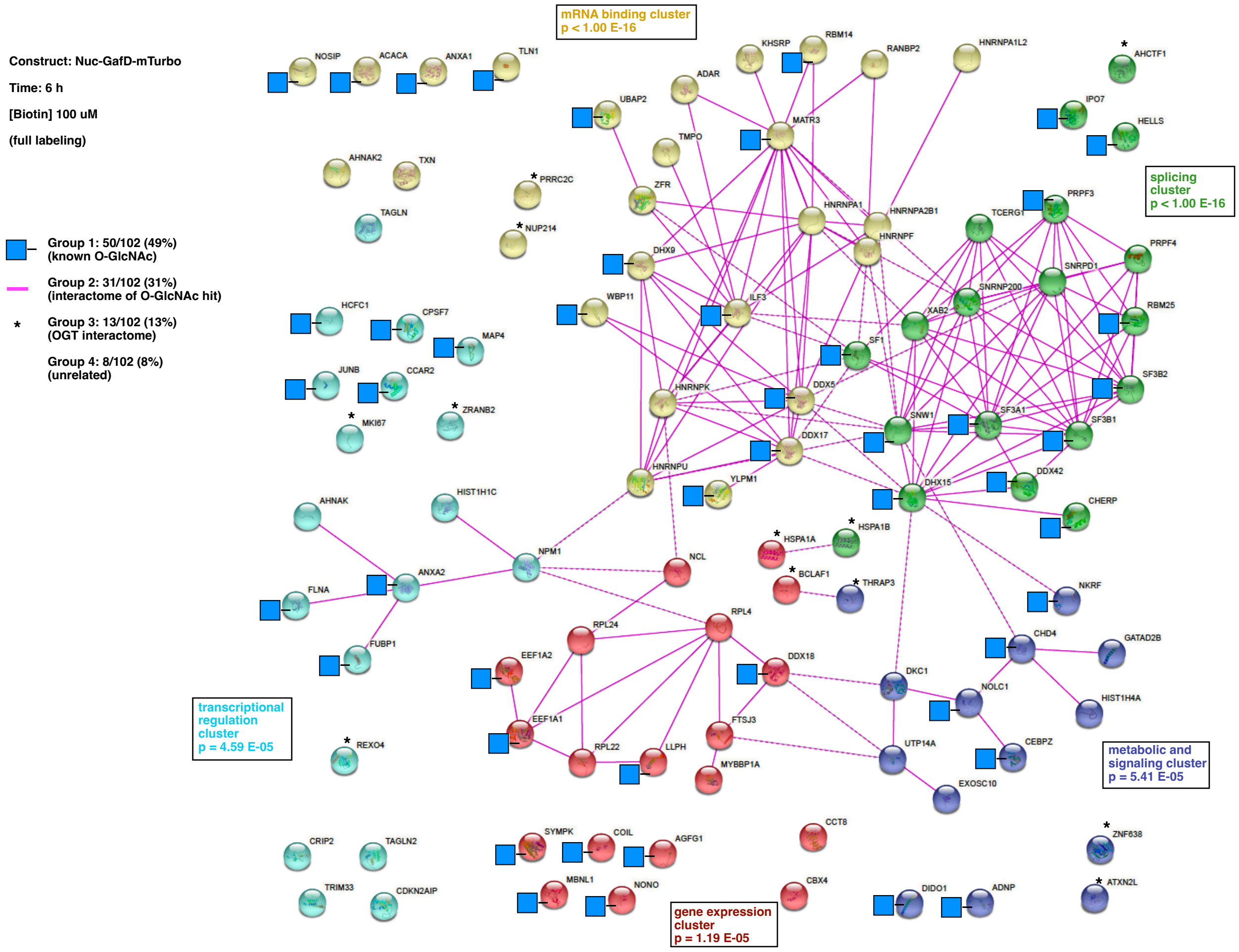


**Supplemental Figure 6:** Interactome analysis between proteins labeled by nuc-GlycoID in the indicated conditions. The 102 hits enriched and exclusive to this condition were input into STRING version 11.5 using the web interface. Analysis was performed as described above. Pink lines represent experimentally-validated protein-protein interactions between targets. The significant gene ontology clusters are labeled with p-values provided in the boxes. The distribution of hits into our arbitrary “Group 1-4” was conducted as labeled in the Figure Legend.

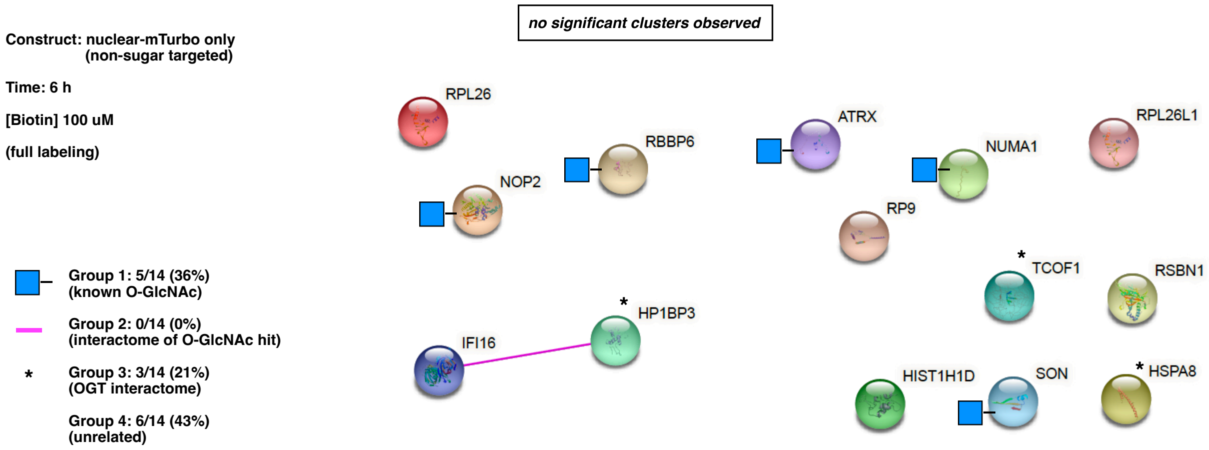


**Supplemental Figure 7:** Interactome analysis between proteins labeled by nuc-mTurbo in the indicated conditions. The 14 hits enriched and exclusive to this condition were input into STRING version 11.5 using the web interface. Analysis was performed as described above. Pink lines represent experimentally-validated protein-protein interactions between targets. The significant gene ontology clusters are labeled with p-values provided in the boxes. The distribution of hits into our arbitrary "Group 1-4" was conducted as labeled in the Figure Legend.

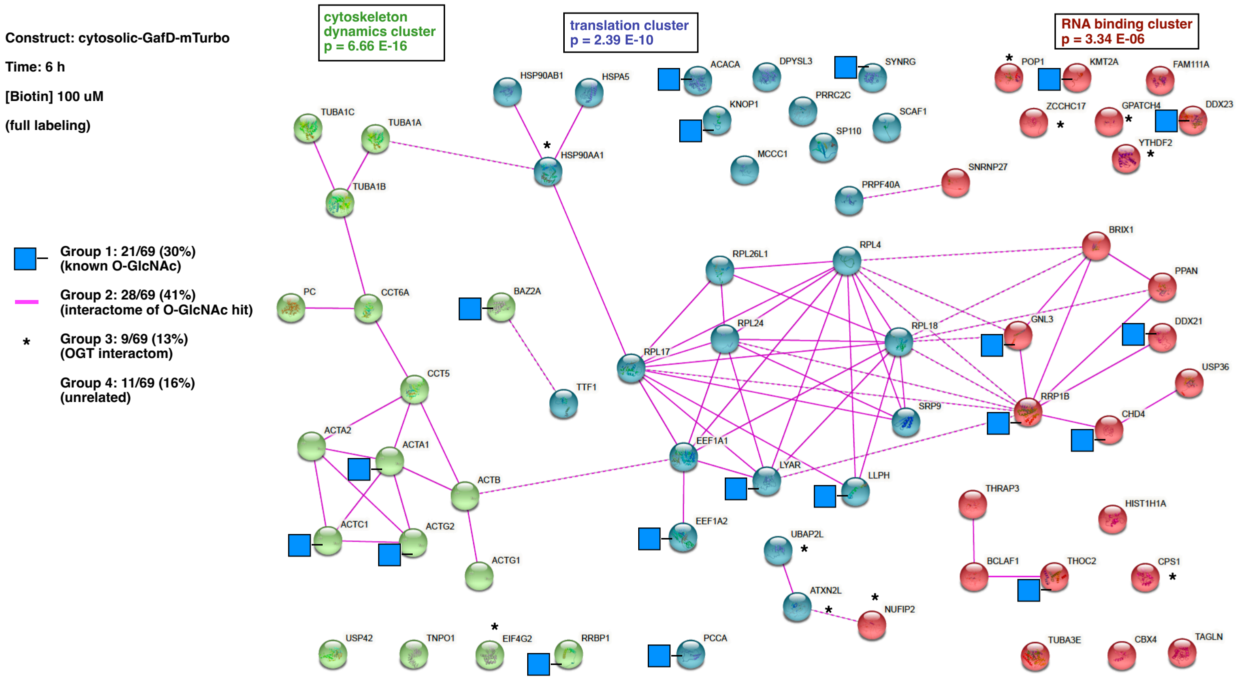


**Supplemental Figure 8:** Interactome analysis between proteins labeled by cyt-GlycoID in the indicated conditions. The 69 hits enriched and exclusive to this condition were input into STRING version 11.5 using the web interface. Analysis was performed as described above. Pink lines represent experimentally-validated protein-protein interactions between targets. The significant gene ontology clusters are labeled with p-values provided in the boxes. The distribution of hits into our arbitrary “Group 1-4” was conducted as labeled in the Figure Legend.

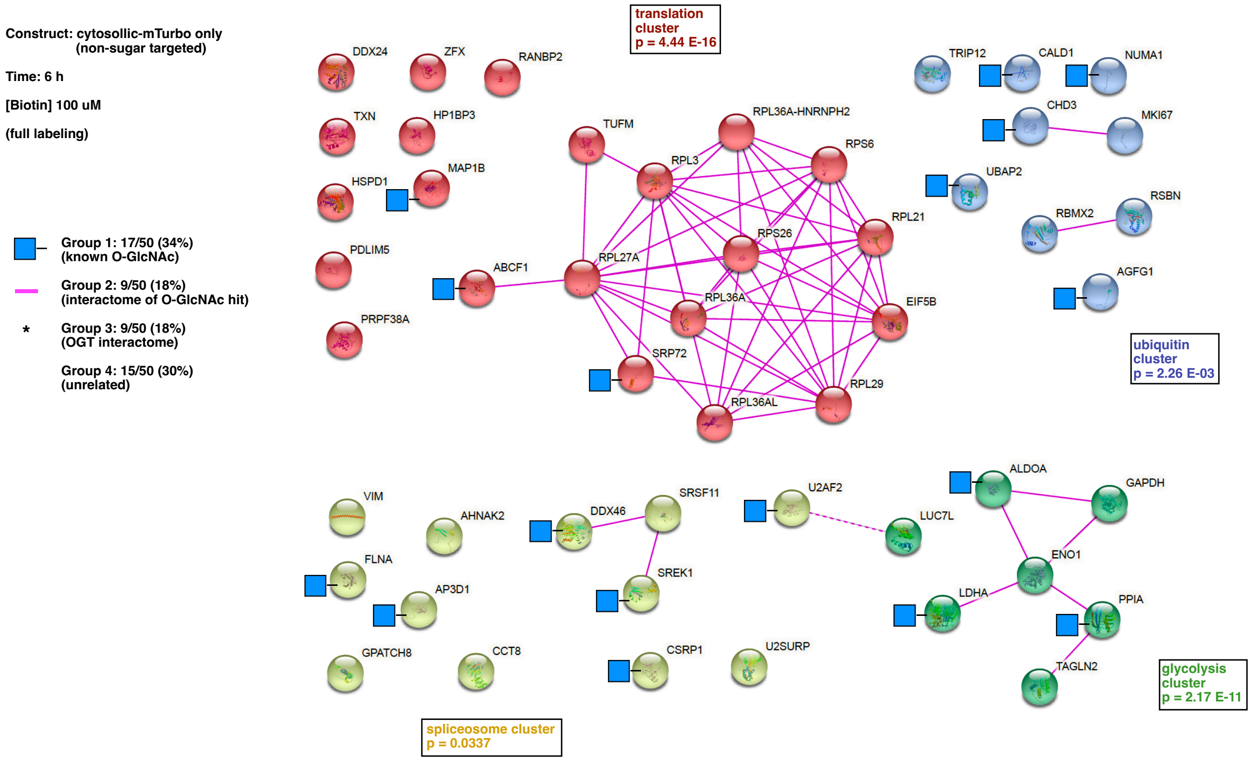


**Supplemental Figure 9:** Interactome analysis between proteins labeled by cyt-mTurbo in the indicated conditions. The 50 hits enriched and exclusive to this condition were input into STRING version 11.5 using the web interface. Analysis was performed as described above. Pink lines represent experimentally-validated protein-protein interactions between targets. The significant gene ontology clusters are labeled with p-values provided in the boxes. The distribution of hits into our arbitrary "Group 1-4" was conducted as labeled in the Figure Legend.

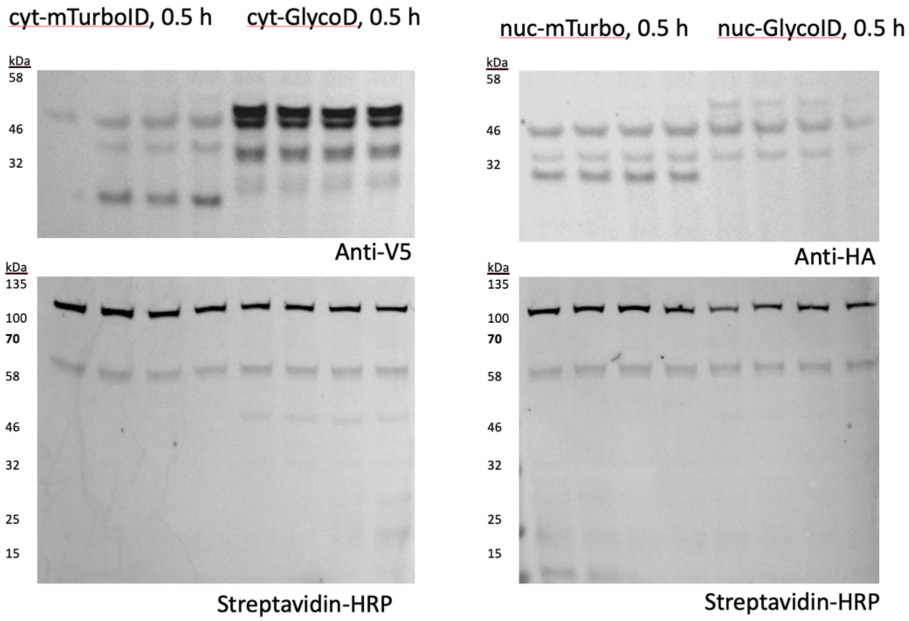


**Supplementary Figure 10A:** Western blot for the expression and activity for cyt-mTurbo, cyt-GlycoID (left blot), nuc-mTurbo, and nuc-GlycoID (right blot) in serum starved HeLa cells used in proteomics experiments. Labeling was conducted for 0.5 hours with 500 μM biotin. Blots were imaged using iBright™ FL1500 instrument.

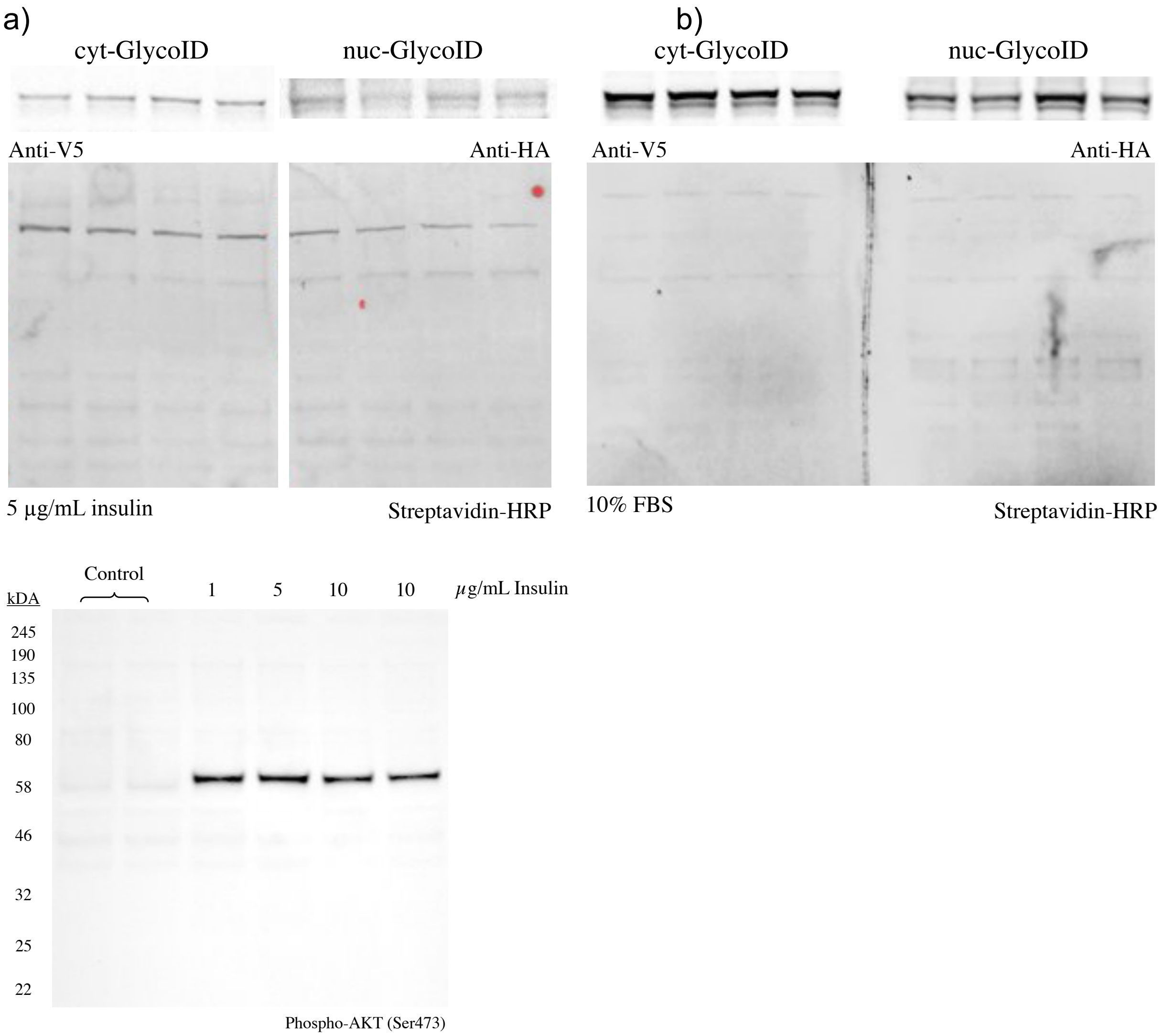


**Supplementary Figure 10B:** Western blots for the expression and activity of cyt-GlycoID and nuc-GlycoID in HeLa cells used in proteomics experiments. a) Shows the labeling activity of cyt/nuc-GlycoID with HeLa cells supplemented with 5 µg/mL insulin with 0.5 hours of labeling with 500 µM biotin. b) Shows the blots of the labeling activity of cyt/nuc-GlycoID with HeLa cells with 10% FBS with 0.5 hours of labeling and 500 µM biotin. C) Shows a blot depicting the effect of insulin on Akt blotted against Phospho-Akt (Ser473) antibody/Anti-Rabbit (1:10,000). Incubating the cells with insulin causes the phosphorylation of Akt.

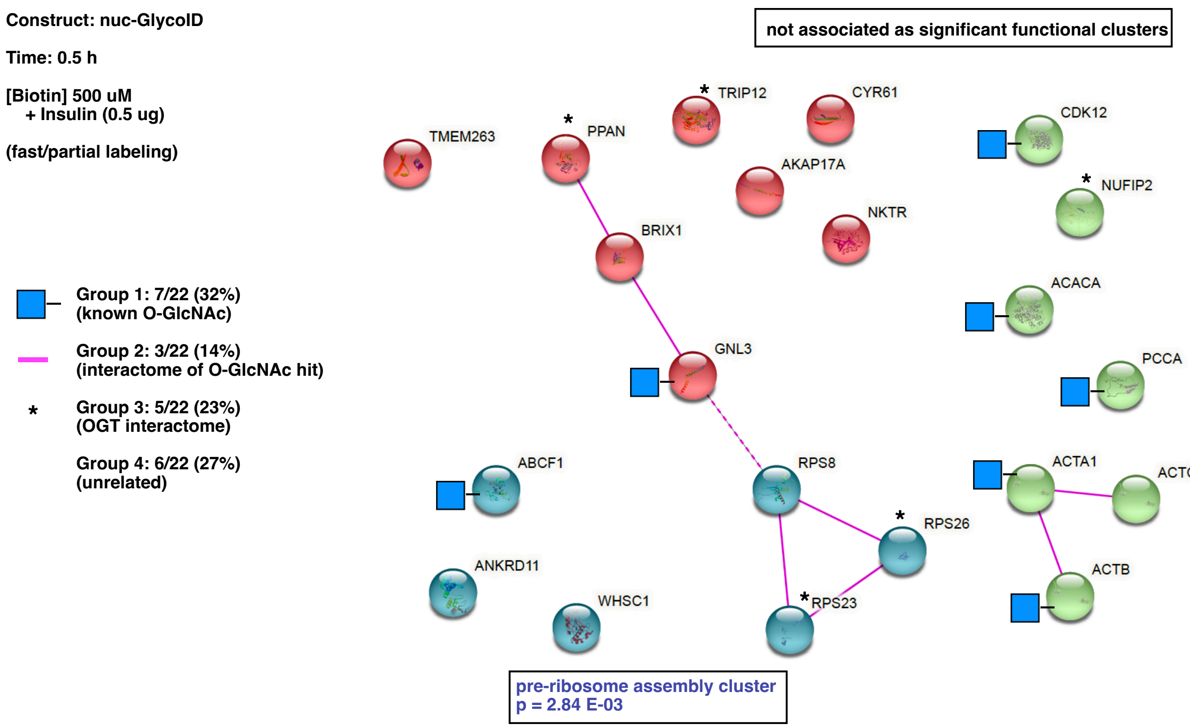


**Supplemental Figure 11:** Interactome analysis between proteins labeled by nuc-GlycoID + insulin in the indicated conditions. The 22 hits enriched and exclusive to this condition were input into STRING version 11.5 using the web interface. Analysis was performed as described above. Pink lines represent experimentally-validated protein-protein interactions between targets. The significant gene ontology clusters are labeled with p-values provided in the boxes. The distribution of hits into our arbitrary "Group 1-4" was conducted as labeled in the Figure Legend.

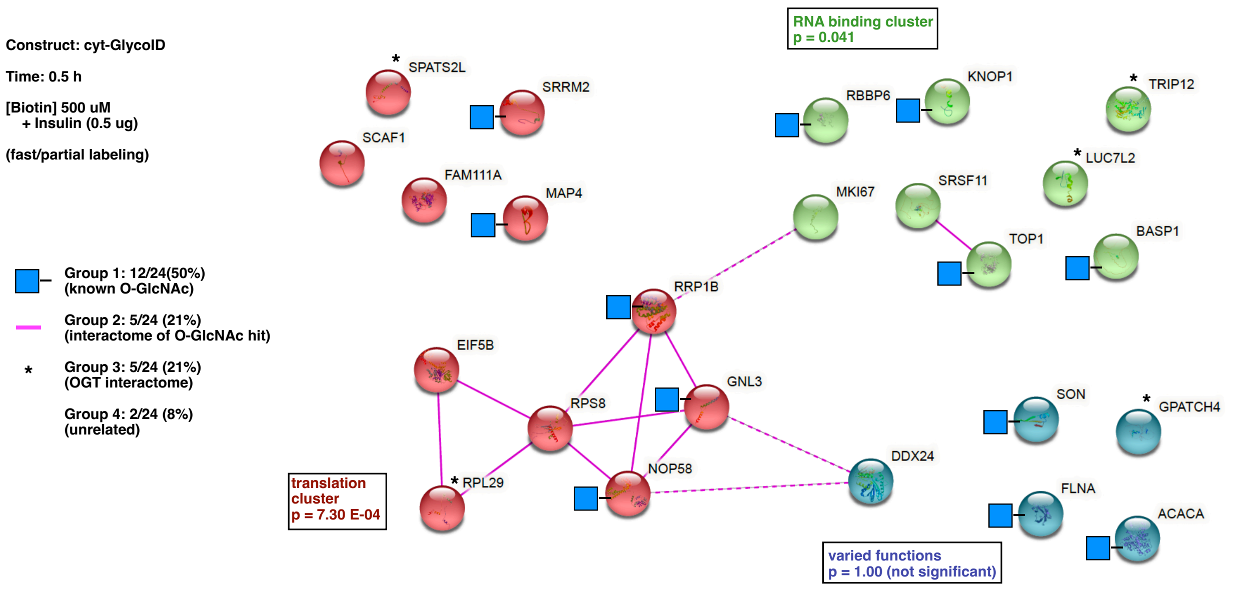


**Supplemental Figure 12:** Interactome analysis between proteins labeled by cyt-GlycoID + insulin in the indicated conditions. The 24 hits enriched and exclusive to this condition were input into STRING version 11.5 using the web interface. Analysis was performed as described above. Pink lines represent experimentally-validated protein-protein interactions between targets. The significant gene ontology clusters are labeled with p-values provided in the boxes. The distribution of hits into our arbitrary “Group 1-4” was conducted as labeled in the Figure Legend.

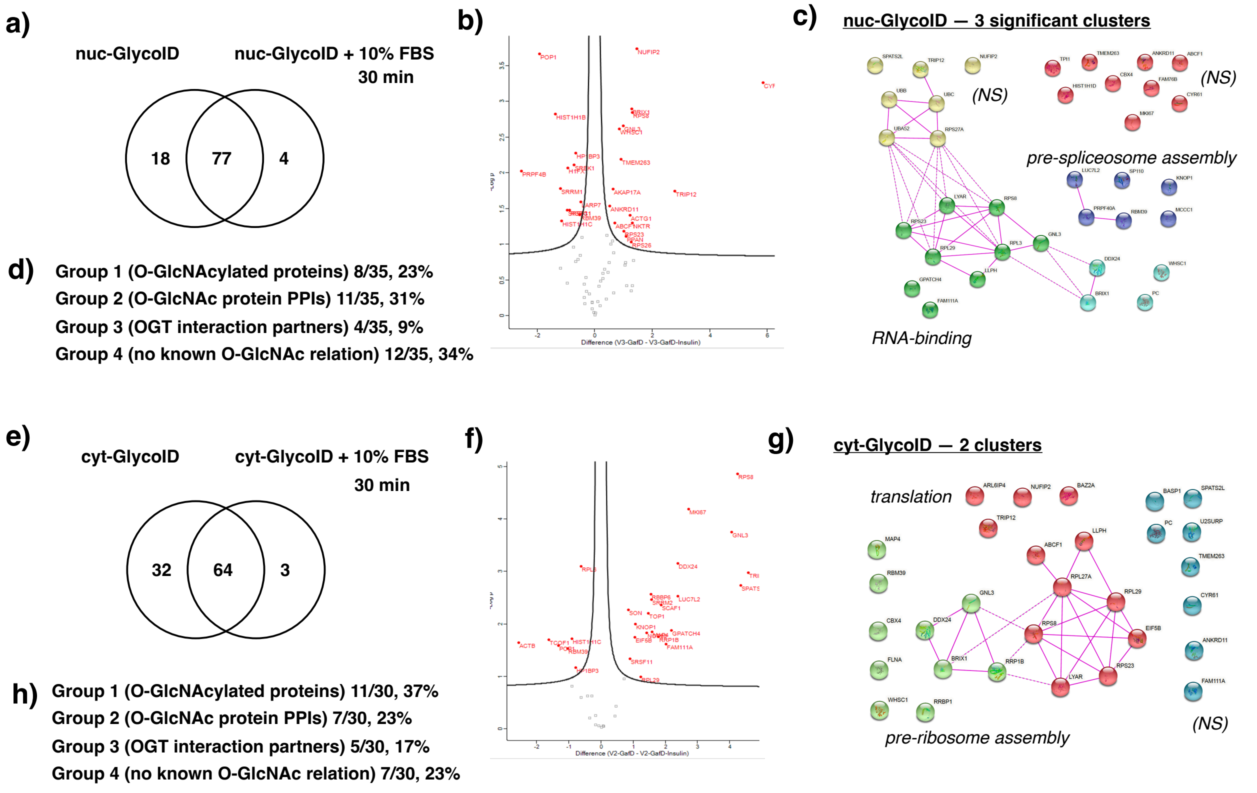


**Supplemental Figure 13:** Analysis of GlycoID constructs between starved and serum fed conditions. ***a***) Exclusive hits between starved nuc-GlycoID vs. 10% fetal bovine serum (FBS)-supplemented nuc-GlycoID. ***b***) Enrichment analysis between starved and fed conditions, statistically significant hits are shown above the volcano plot cutoffs. ***c***) Physical interactions between serum-fed hits reveal functional clusters with key O-GlcNAc linkages. ***d***) The protein groups labeled by nuc-GlycoID serum-fed hits, as defined in the main text. ***e***)-***h***) *A*nalysis for starved cyt-GlycoID vs. 10% FBS-supplemented cyt-GlycoID. Full-sized STRING plots with labeled O-GlcNAc hits are found in **Supporting Figures 14-15**.

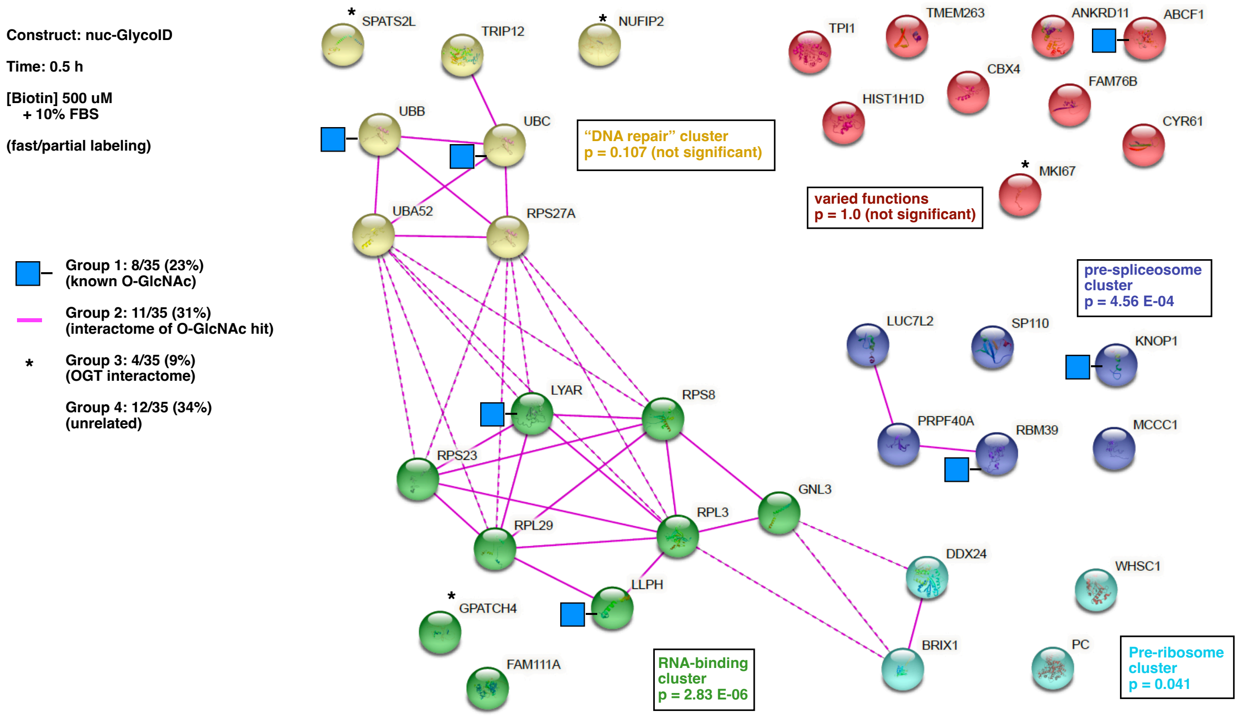


**Supplemental Figure 14:** Interactome analysis between proteins labeled by nuc-GlycoID + insulin in the indicated conditions. The 35 hits enriched and exclusive to this condition were input into STRING version 11.5 using the web interface. Analysis was performed as described above. Pink lines represent experimentally-validated protein-protein interactions between targets. The significant gene ontology clusters are labeled with p-values provided in the boxes. The distribution of hits into our arbitrary "Group 1-4" was conducted as labeled in the Figure Legend.

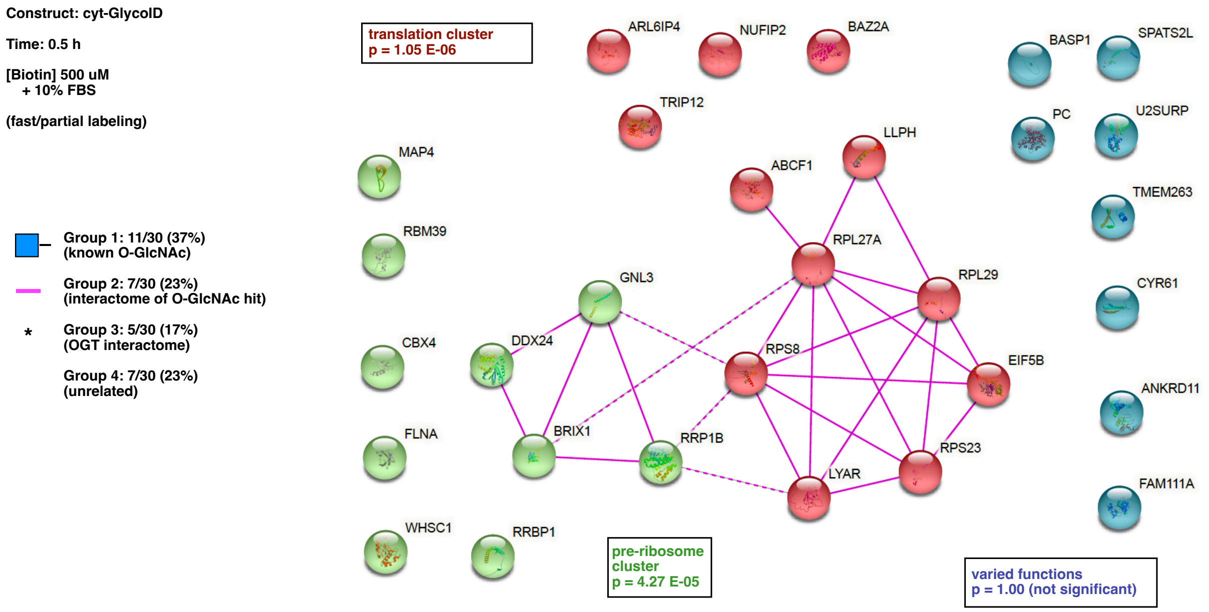


**Supplemental Figure 15:** Interactome analysis between proteins labeled by nuc-GlycoID + insulin in the indicated conditions. The 30 hits enriched and exclusive to this condition were input into STRING version 11.5 using the web interface. Analysis was performed as described above. Pink lines represent experimentally-validated protein-protein interactions between targets. The significant gene ontology clusters are labeled with p-values provided in the boxes. The distribution of hits into our arbitrary "Group 1-4" was conducted as labeled in the Figure Legend.
